## Supplementary material for "Closely related, yet phenotypically different - Genome assemblies of two sister species of widow spiders: *Latrodectus hasselti* and *L. katipo*, Theridiidae": SM

**Supplemetary material.**

Contents

Specimens’ metadata	2

Genome assembly	3

Genome annotation	6

Repetitive elements	11

Programs and commands	14

### Specimens’ metadata

Table S1. Specimens’ metadata.

| **ID number** | **Species** | **n** | **Type of sequencing** | **Sequencing platform** | **Sex** | **Family origin** | | | |
| --- | --- | --- | --- | --- | --- | --- | --- | --- | --- |
|  |  |  |  |  |  | **Country** | **Locality** | **Latitude** | **Longitude** |
| 64K123 | *Latrodectus katipo* | 1 | Long reads, CLR; Short reads for polishing | PacBio Sequel II; Illumina NovaSeq X Plus (PE150) | Female | New Zealand | Kaitorete Spit | -43.827880 | 172.669921 |
| 60K233 | *Latrodectus katipo* | 1 | Short reads, Hi-C | Illumina Novaseq 6000 (PE150) | Female | New Zealand | Kaitorete Spit | -43.827880 | 172.669921 |
| KPOOL3 | *Latrodectus katipo* | 9 | RNA pool short-read, annotation | Illumina NovaSeq X Plus (PE150) | Male | Germany | University of Hamburg | NA | NA |
| KPOOL4 | *Latrodectus katipo* | 9 | RNA pool short-read, annotation | Illumina NovaSeq X Plus (PE150) | Male | Germany | University of Hamburg | NA | NA |
| KK5076 | *Latrodectus katipo* | 1 | Whole genome, short reads | Illumina NovaSeq X Plus (PE150) | Male | Germany | University of Hamburg | NA | NA |
| KKKK5621 | *Latrodectus katipo* | 1 | Whole genome, short reads | Illumina NovaSeq X Plus (PE150) | Male | Germany | University of Hamburg | NA | NA |
| 49H2389 | *Latrodectus hasselti* | 1 | Long reads, CCS | PacBio Revio | Female | New Zealand | Alexandra | -45.254600 | 169.402691 |
| HPOOL1 | *Latrodectus hasselti* | 9 | RNA pool short-read, annotation | Illumina NovaSeq X Plus (PE150) | Male | Germany | University of Hamburg | NA | NA |
| HPOOL2 | *Latrodectus hasselti* | 9 | RNA pool short-read, annotation | Illumina NovaSeq X Plus (PE150) | Male | Germany | University of Hamburg | NA | NA |
| AFRB13 | *Latrodectus hasselti* | 1 | RNA, short reads | Illumina HiSeq 2500 | Female | Canada | University of Toronto | NA | NA |
| AMRB25 | *Latrodectus hasselti* | 1 | RNA, short reads | Illumina HiSeq 2500 | Female | Canada | University of Toronto | NA | NA |
| PFRB15 | *Latrodectus hasselti* | 1 | RNA, short reads | Illumina HiSeq 2500 | Female | Canada | University of Toronto | NA | NA |
| PMSRB53 | *Latrodectus hasselti* | 1 | RNA, short reads | Illumina HiSeq 2500 | Female | Canada | University of Toronto | NA | NA |

### Genome assembly

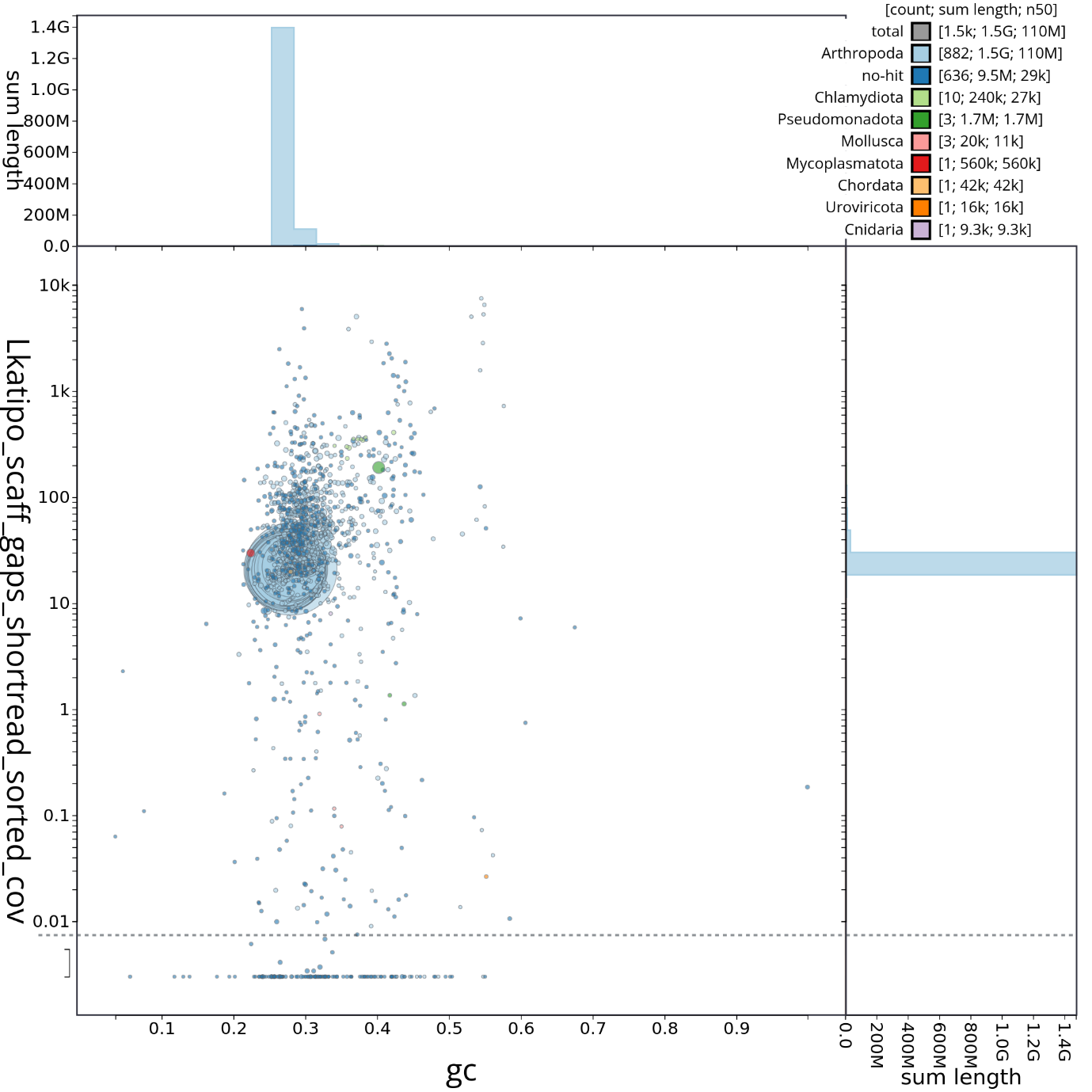

Figure S1. Blobtools plot of contamination in *L. katipo* assembly.

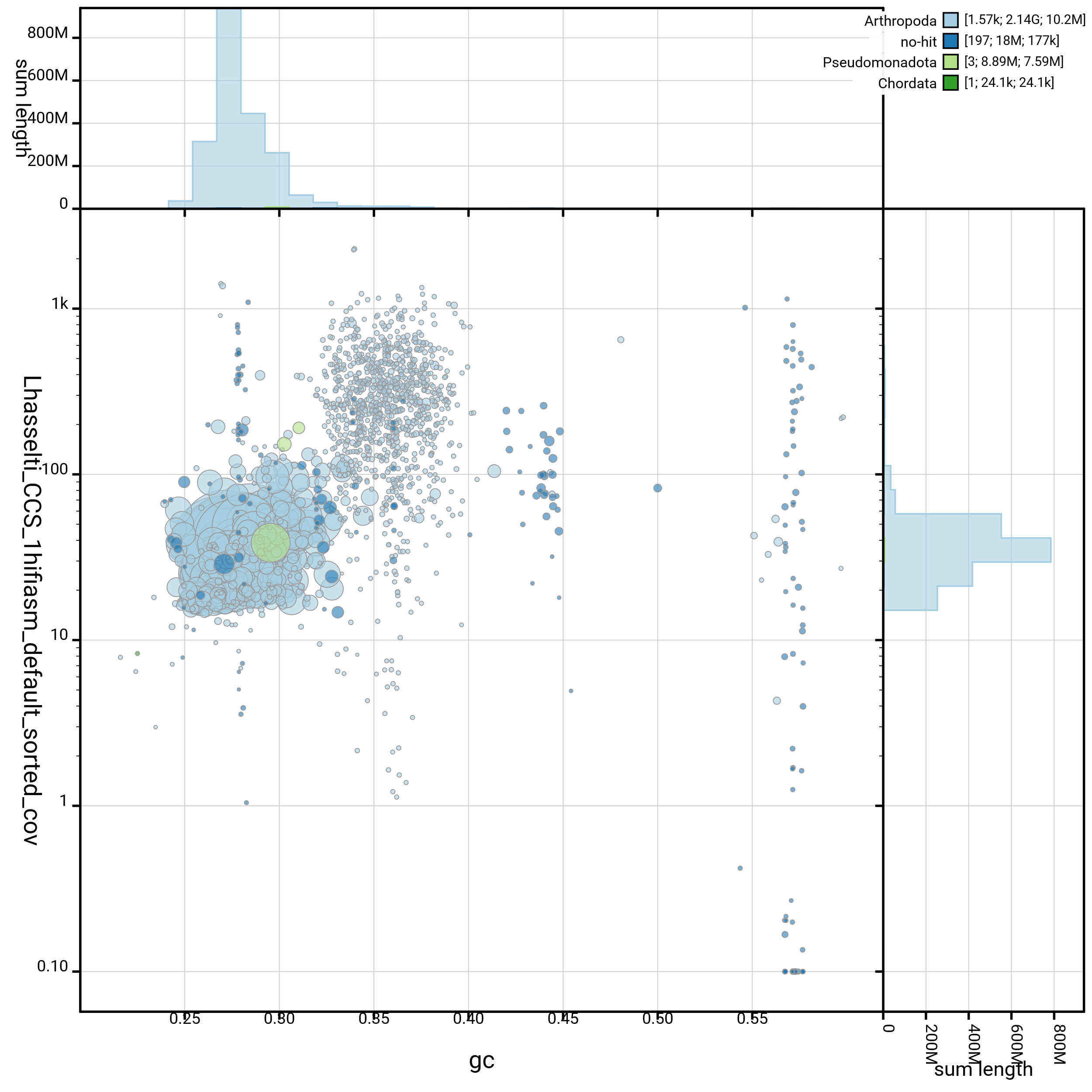

Figure S2. Blobtools plot of contamination in *L. hasselti* assembly.

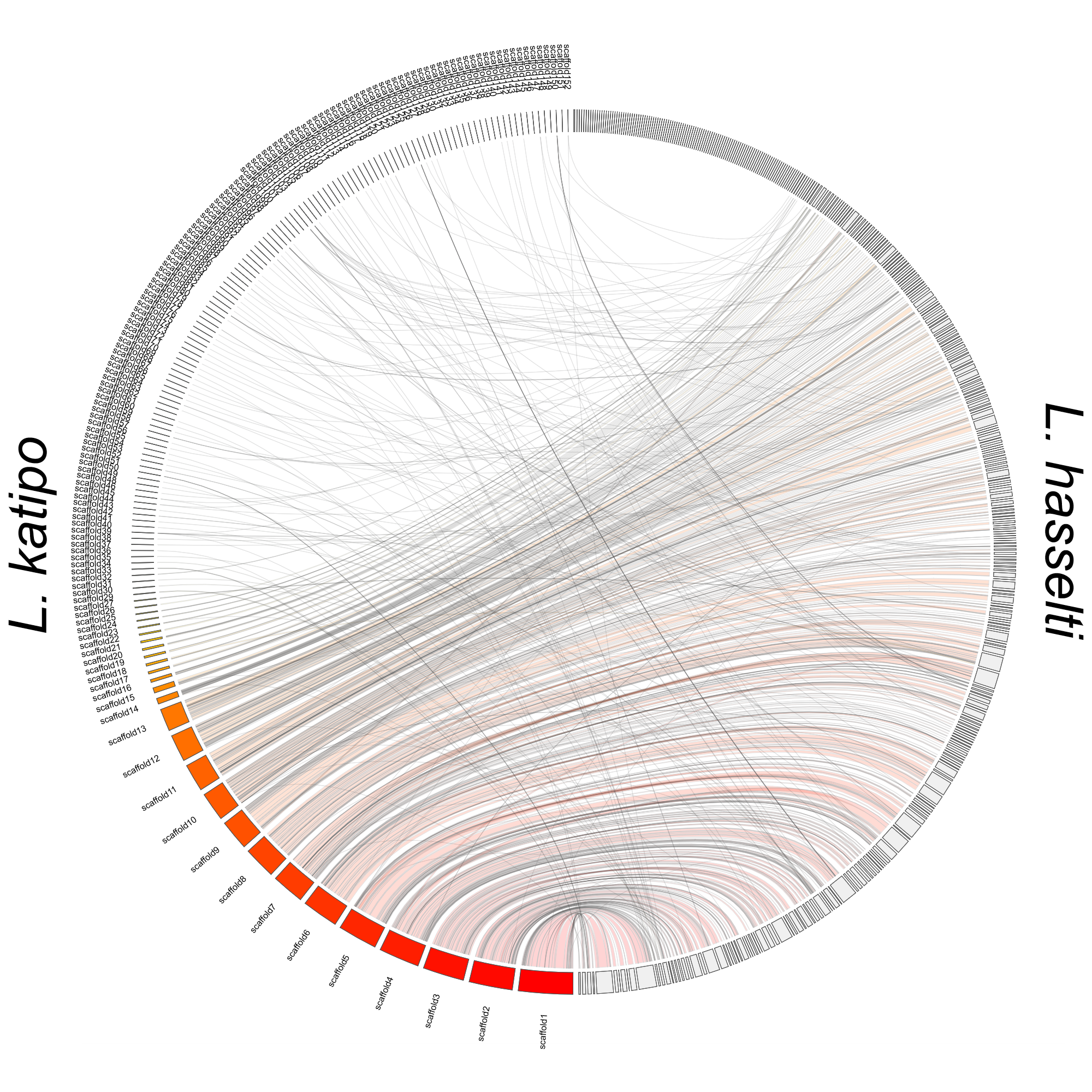

Figure S3. Jupiter plot of *L. hasselti* assembly mapped against *L. katipo* assembly. All scaffolds and contigs from both final assemblies were used as an input.

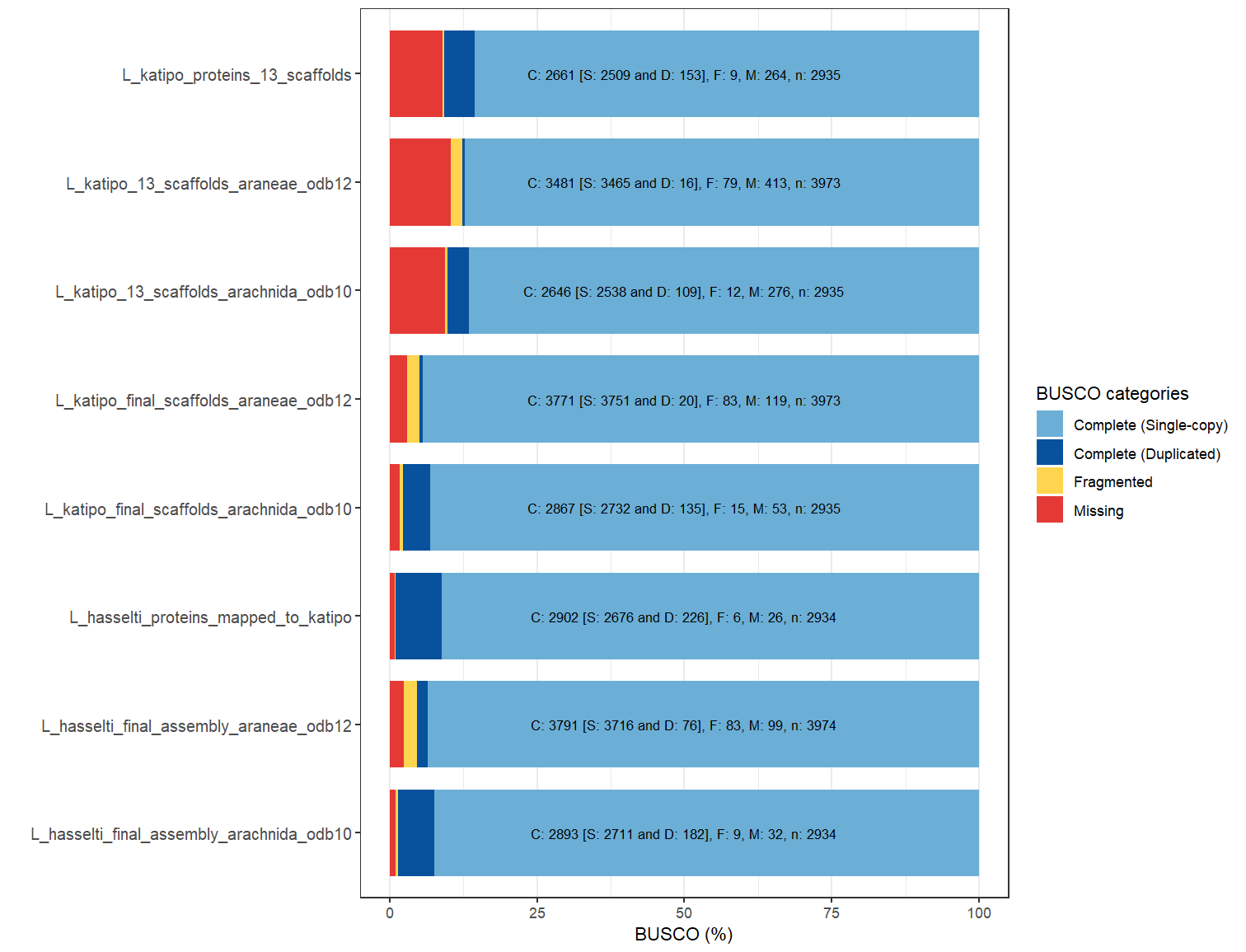

Figure S4. BUSCO scores for genome assemblies and protein sequences.

Table S2. BUSCO scores for genome assemblies and protein sequences. C – complete, S – single-copy, D – duplicated, F -fragmented, M – missing.

| **Assmelby** | **C_%** | **C_n_genes** | **S_%** | **S_n_genes** | **D_%** | **D_n_genes** | **F_%** | **F_n_genes** | **M_%** | **M_n_genes** | **Total_n_genes** |
| --- | --- | --- | --- | --- | --- | --- | --- | --- | --- | --- | --- |
| L. hasselti final assembly arachnida_odb 10 | 98.6 | 2893 | 92.4 | 2711 | 6.2 | 182 | 0.3 | 9 | 1.1 | 32 | 2934 |
| L. hasselti final assembly araneae_odb 12 | 95.4 | 3791 | 93.5 | 3716 | 1.9 | 76 | 2.1 | 83 | 2.5 | 99 | 3974 |
| L. hasselti protein sequences from contigs mapped to L. katipo genome arachnida_odb 10 | 98.9 | 2902 | 91.2 | 2676 | 7.7 | 226 | 0.2 | 6 | 0.9 | 26 | 2934 |
| L. katipo final assemly all scaffolds arachnida_odb 10 | 97.7 | 2867 | 93.1 | 2732 | 4.6 | 135 | 0.5 | 15 | 1.8 | 53 | 2934 |
| L. katipo final assemly all scaffolds araneae_odb 12 | 94.9 | 3771 | 94.4 | 3751 | 0.5 | 20 | 2.1 | 83 | 3 | 119 | 3974 |
| L. katipo 13 scaffolds arachnida_odb 10 | 90.2 | 2646 | 86.5 | 2538 | 3.7 | 109 | 0.4 | 12 | 9.4 | 276 | 2934 |
| L. katipo 13 scaffolds araneae_odb 12 | 87.6 | 3481 | 87.2 | 3465 | 0.4 | 16 | 2 | 79 | 10.4 | 413 | 3974 |
| L. katipo protein sequences 13 scaffolds arachnida_odb 10 | 90.7 | 2661 | 85.5 | 2509 | 5.2 | 153 | 0.3 | 9 | 9 | 264 | 2934 |

### Genome annotation

Table S3. Total COG counts.

| **General COG** | **Specific COG** | **COG letter** | ***L. hasselti*, n** | ***L. katipo* n** | ***L. hasselti*, %** | ***L. katipo*, %** |
| --- | --- | --- | --- | --- | --- | --- |
| Information storage and processing | Translation, ribosomal structure and biogenesis | J | 415 | 370 | 3.88 | 4.05 |
| Information storage and processing | RNA processing and modification | A | 446 | 389 | 4.17 | 4.26 |
| Information storage and processing | Transcription | K | 861 | 712 | 8.04 | 7.79 |
| Information storage and processing | Replication, recombination and repair | L | 303 | 253 | 2.83 | 2.77 |
| Information storage and processing | Chromatin structure and dynamics | B | 151 | 128 | 1.41 | 1.40 |
| Cellular processes and signalling | Cell cycle control, cell division, chromosome partitioning | D | 287 | 230 | 2.68 | 2.52 |
| Cellular processes and signalling | Nuclear structure | Y | 34 | 26 | 0.32 | 0.28 |
| Cellular processes and signalling | Defense mechanisms | V | 85 | 72 | 0.79 | 0.79 |
| Cellular processes and signalling | Signal transduction mechanisms | T | 1525 | 1258 | 14.25 | 13.76 |
| Cellular processes and signalling | Cell wall/membrane/envelope biogenesis | M | 86 | 74 | 0.80 | 0.81 |
| Cellular processes and signalling | Cell motility | N | 11 | 8 | 0.10 | 0.09 |
| Cellular processes and signalling | Cytoskeleton | Z | 335 | 279 | 3.13 | 3.05 |
| Cellular processes and signalling | Extracellular structures | W | 76 | 56 | 0.71 | 0.61 |
| Cellular processes and signalling | Intracellular trafficking, secretion, and vesicular transport | U | 566 | 491 | 5.29 | 5.37 |
| Cellular processes and signalling | Posttranslational modification, protein turnover, chaperones | O | 1029 | 876 | 9.61 | 9.58 |
| Metabolism | Energy production and conversion | C | 271 | 240 | 2.53 | 2.63 |
| Metabolism | Carbohydrate transport and metabolism | G | 339 | 305 | 3.17 | 3.34 |
| Metabolism | Amino acid transport and metabolism | E | 280 | 257 | 2.62 | 2.81 |
| Metabolism | Nucleotide transport and metabolism | F | 115 | 103 | 1.07 | 1.13 |
| Metabolism | Coenzyme transport and metabolism | H | 116 | 100 | 1.08 | 1.09 |
| Metabolism | Lipid transport and metabolism | I | 426 | 375 | 3.98 | 4.10 |
| Metabolism | Inorganic ion transport and metabolism | P | 436 | 363 | 4.07 | 3.97 |
| Metabolism | Secondary metabolites biosynthesis, transport and catabolism | Q | 279 | 240 | 2.61 | 2.63 |
| Poorly characterized | General function prediction only | R | 0 | 0 | 0.00 | 0.00 |
| Poorly characterized | Function unknown | S | 2231 | 1935 | 20.84 | 21.17 |
| Total |  |  | 10703 | 9140 | 100.00 | 100.00 |

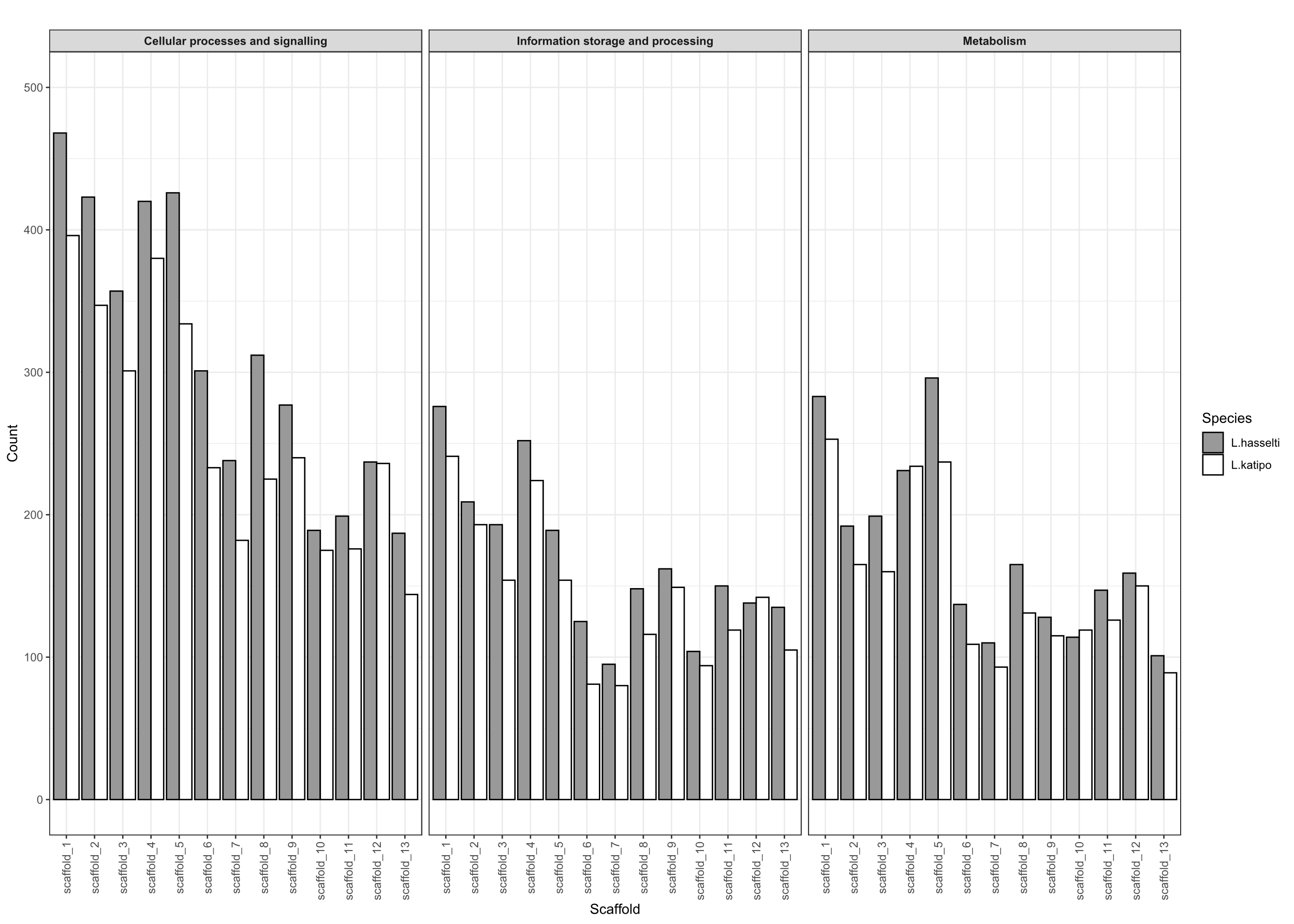

Fig. S5. Distribution of general COG counts across scaffolds.

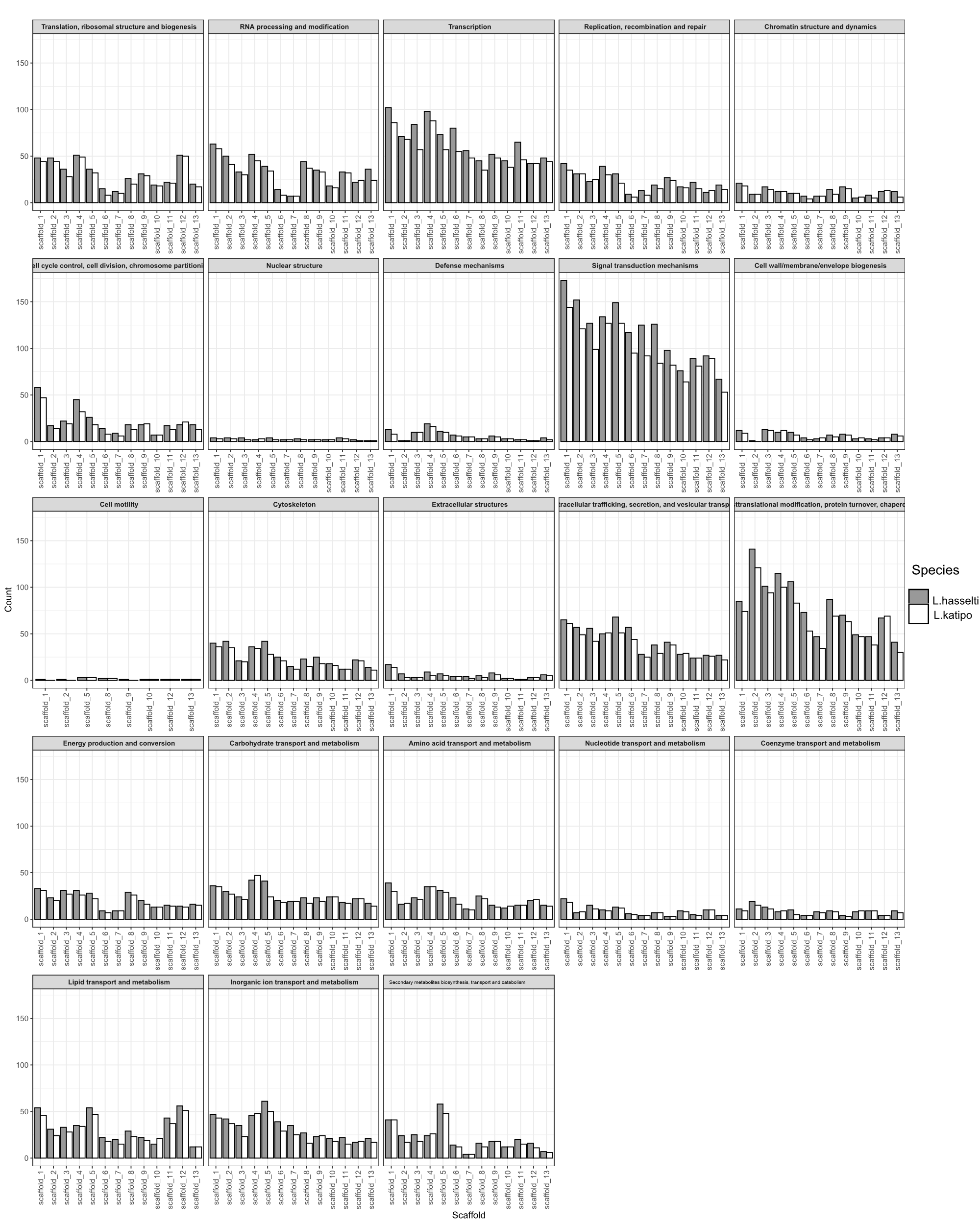

Fig. S6. Distribution of specific COG counts across scaffolds

Table S4. Orthofinder statistics per species.

|  | *Argiope bruennichi* | *Latrodectus elegans* | *Latrodectus hasselti* | *Latrodectus hesperus* | *Latrodectus katipo* | *Nephila pilipes* | *Oedothorax gibbosus* | *Parasteatoda tepidariorum* | *Pimoa clavata* | *Stegodyphus dumicola* | *Tetragnatha kauaiensis* |
| --- | --- | --- | --- | --- | --- | --- | --- | --- | --- | --- | --- |
| Number of genes | 30141 | 20167 | 15111 | 17364 | 12706 | 72441 | 32682 | 34165 | 30370 | 29836 | 42723 |
| Number of genes in orthogroups | 29174 | 16631 | 14840 | 14794 | 12469 | 54215 | 26168 | 33171 | 29164 | 28566 | 31897 |
| Number of unassigned genes | 967 | 3536 | 271 | 2570 | 237 | 18226 | 6514 | 994 | 1206 | 1270 | 10826 |
| Percentage of genes in orthogroups | 96.8 | 82.5 | 98.2 | 85.2 | 98.1 | 74.8 | 80.1 | 97.1 | 96 | 95.7 | 74.7 |
| Percentage of unassigned genes | 3.2 | 17.5 | 1.8 | 14.8 | 1.9 | 25.2 | 19.9 | 2.9 | 4 | 4.3 | 25.3 |
| Number of orthogroups containing species | 12706 | 10064 | 12246 | 9213 | 10670 | 16888 | 13522 | 12689 | 12654 | 13974 | 14594 |
| Percentage of orthogroups containing species | 53.6 | 42.5 | 51.7 | 38.9 | 45 | 71.3 | 57.1 | 53.6 | 53.4 | 59 | 61.6 |
| Number of species-specific orthogroups | 365 | 442 | 6 | 74 | 2 | 2412 | 653 | 473 | 306 | 352 | 1027 |
| Number of genes in species-specific orthogroups | 2031 | 1913 | 22 | 183 | 4 | 13784 | 3014 | 2334 | 1093 | 1252 | 4502 |
| Percentage of genes in species-specific orthogroups | 6.7 | 9.5 | 0.1 | 1.1 | 0 | 19 | 9.2 | 6.8 | 3.6 | 4.2 | 10.5 |

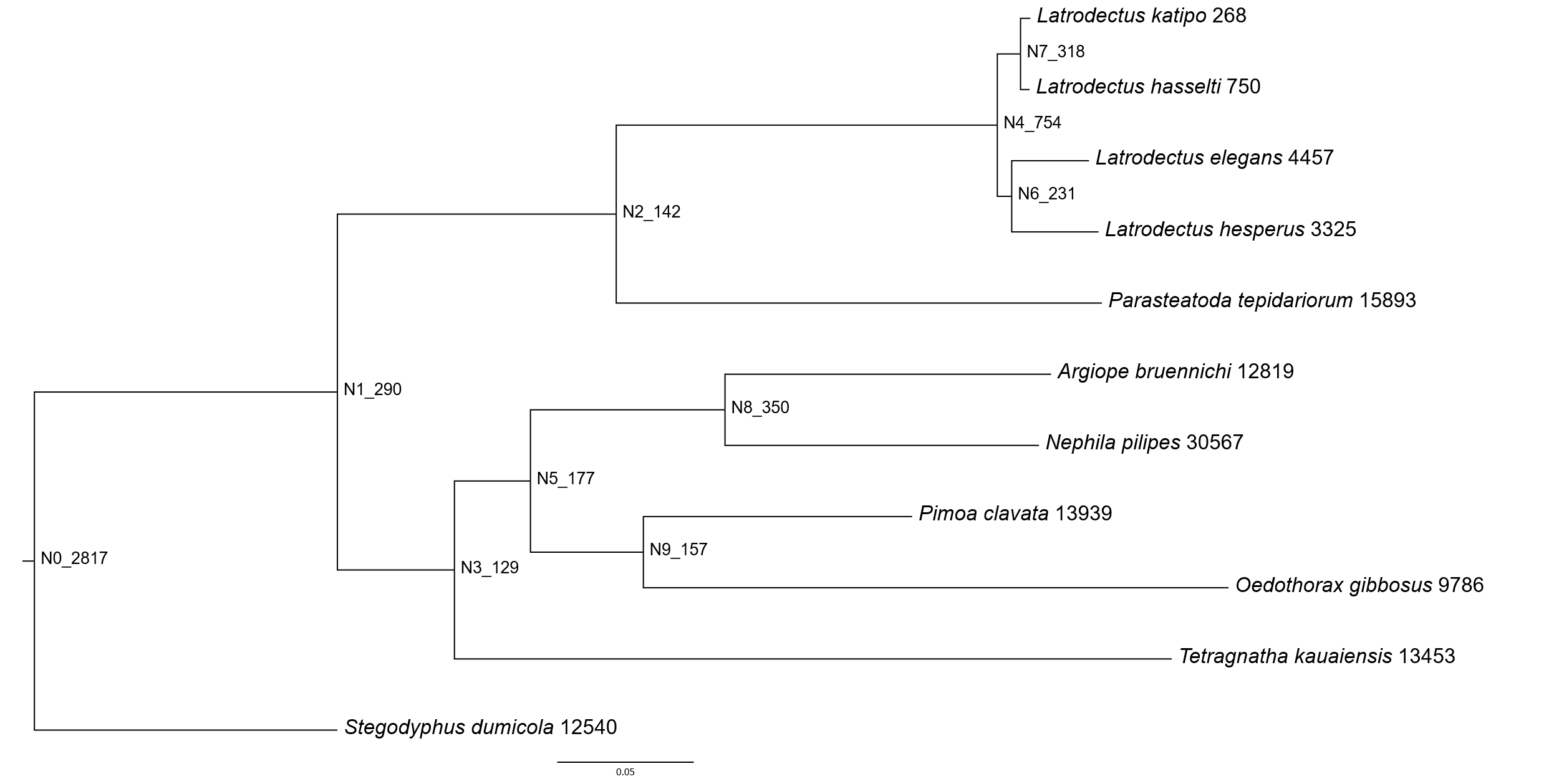

Figure S7. Combined species tree based on sequences predicted by annotation. Numbers at tips and nodes (numbered N0 to N9) represent number of duplicated genes.

### Repetitive elements

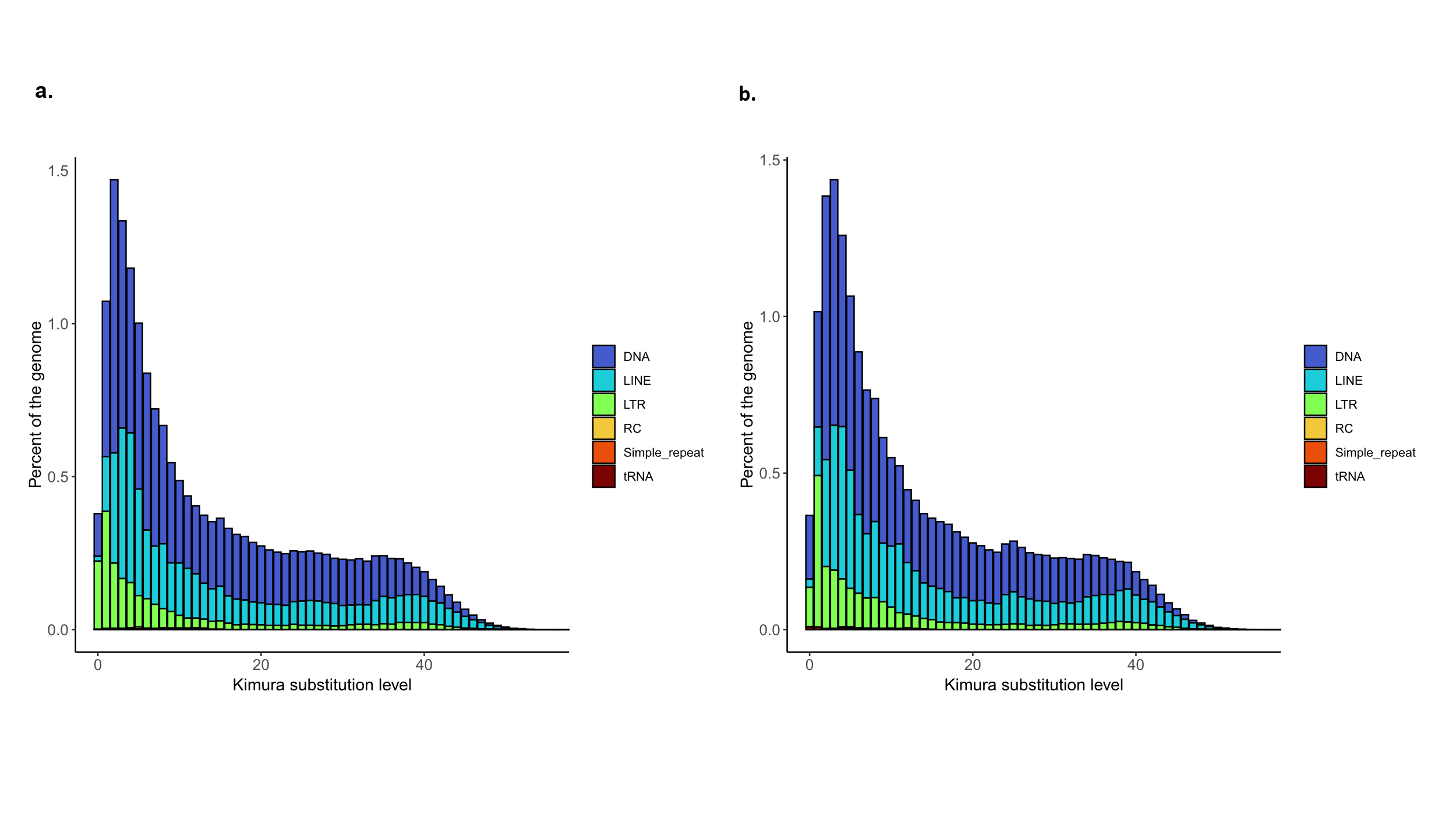

Figure S8. Repetitive elements landscapes showing only classified repeats. a. *L. katipo*; b. *L. hasselti*.

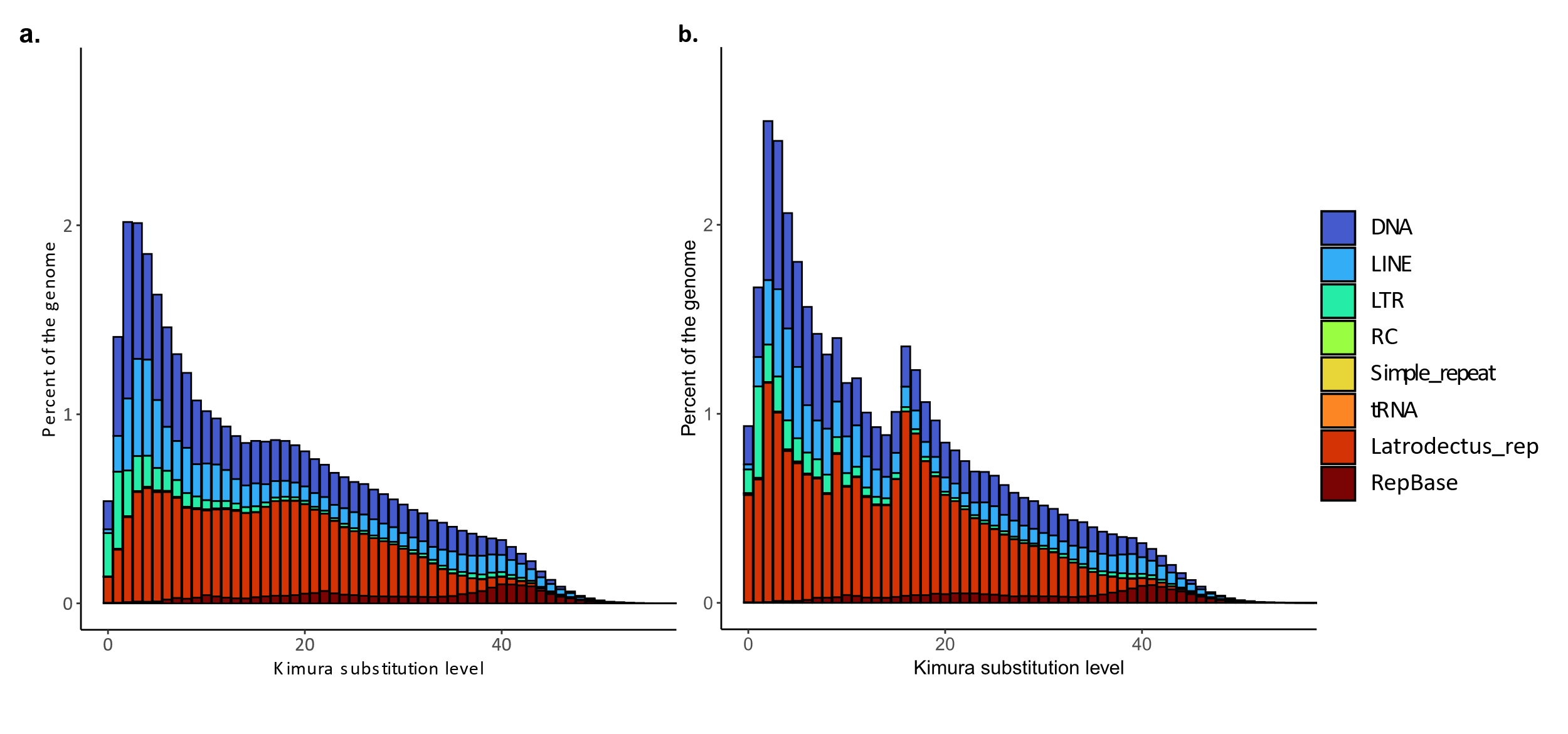

Figure S9. Repetitive elements landscapes showing all repeats. a. *L. katipo*; b. *L. hasselti*. Latrodectus_rep are repetitive elements identified by RepeatModeler but absent in RepBase. RepBase are repetitive elements present in RepBase but not assigned to any class of repeats.

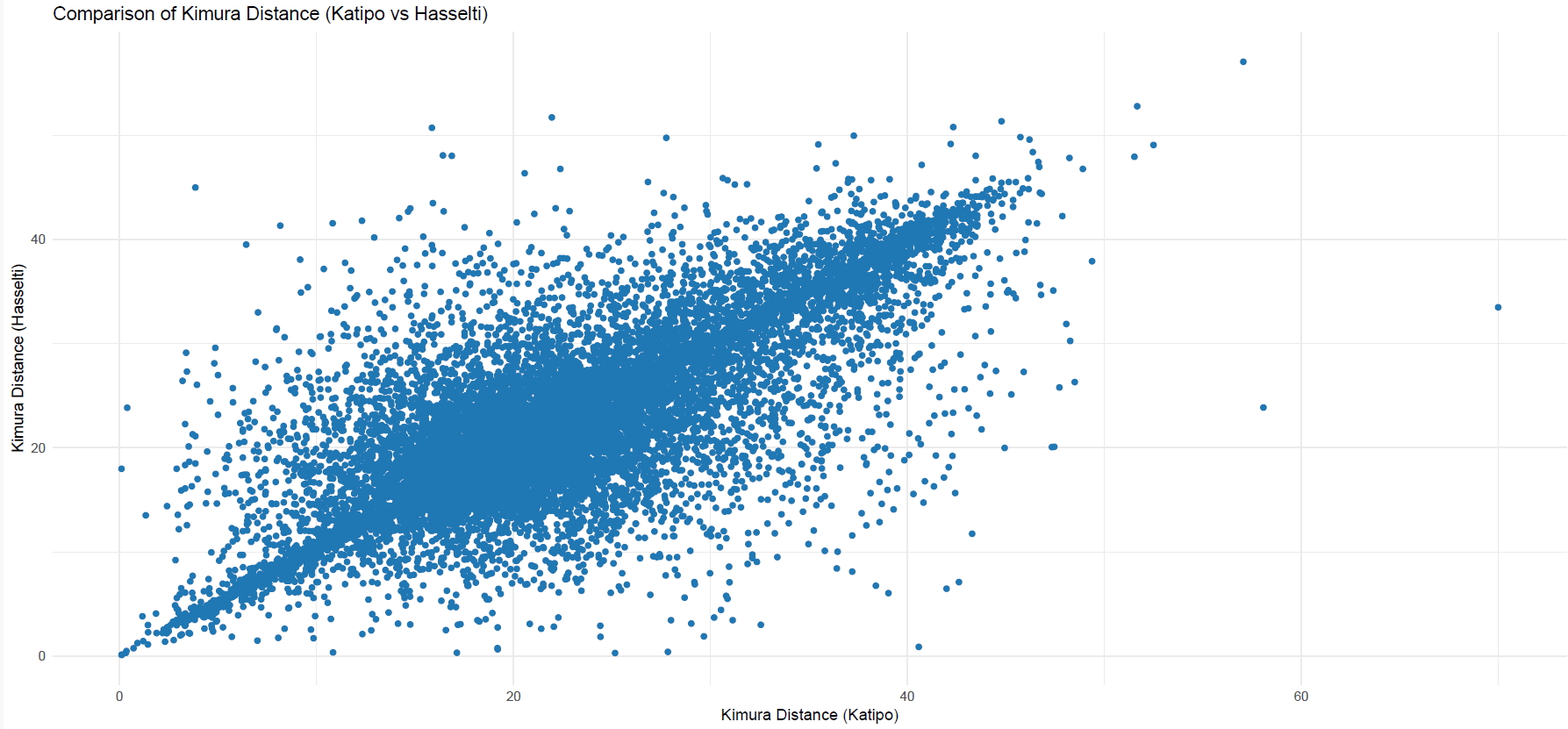

Figure S10. Kimura distances of the same repetitive elements plotted against each other. x-axis *L. katipo*, y-axis *L. hasselti*. The t-test suggests that the sets of Kimura distance in the shared repeats are not significantly different (t = -0.70019, df = 30936, p-value = 0.4838, *L. katipo* mean Kimura distance= 22.08074, *L. hasselti* mean Kimura distance=22.14623).

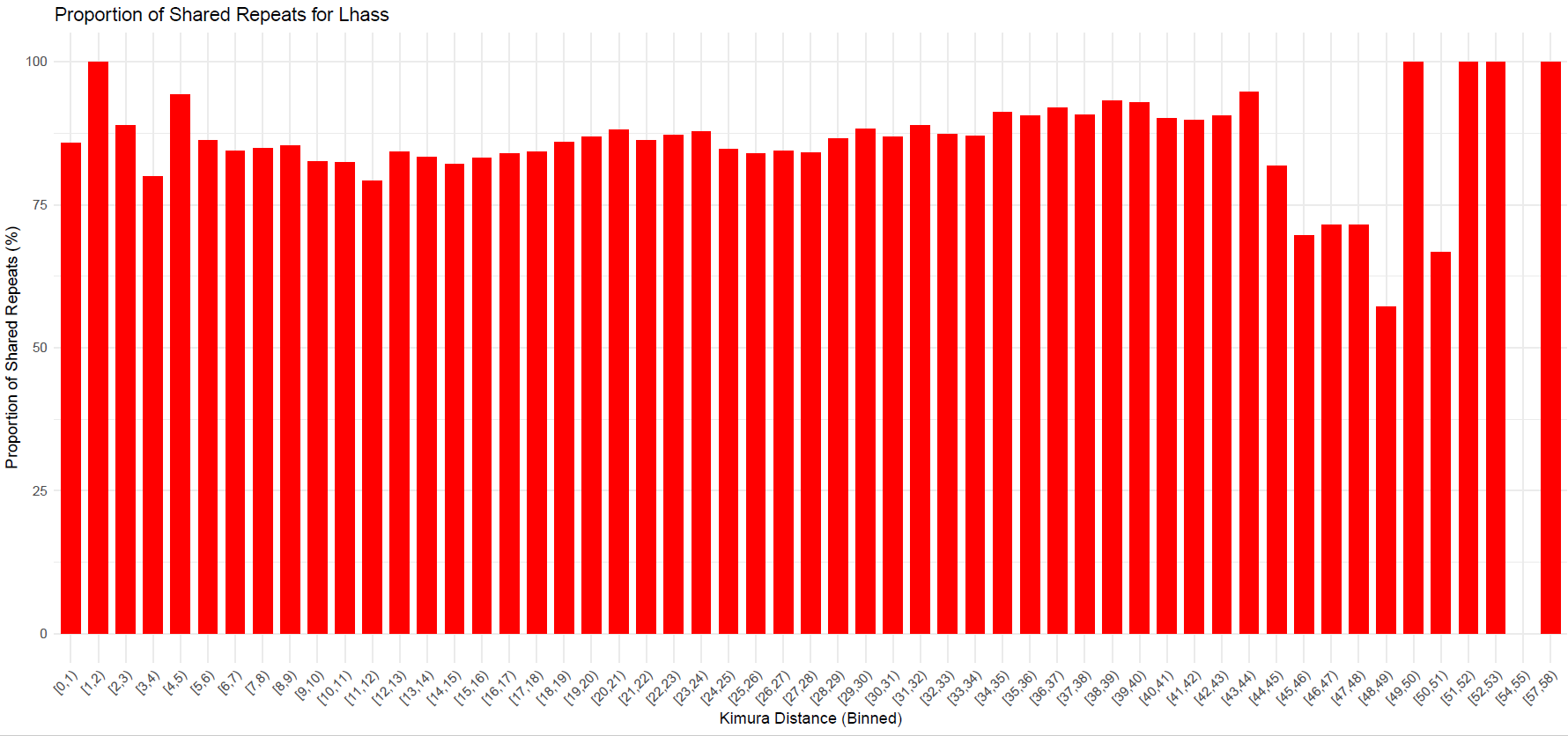

Figure S11. Proportion of shared repeats between the two species in total number of repeats in *L. hasselti* arranged by Kimura distance. One bar represents 1% of the maximum Kimura distance.

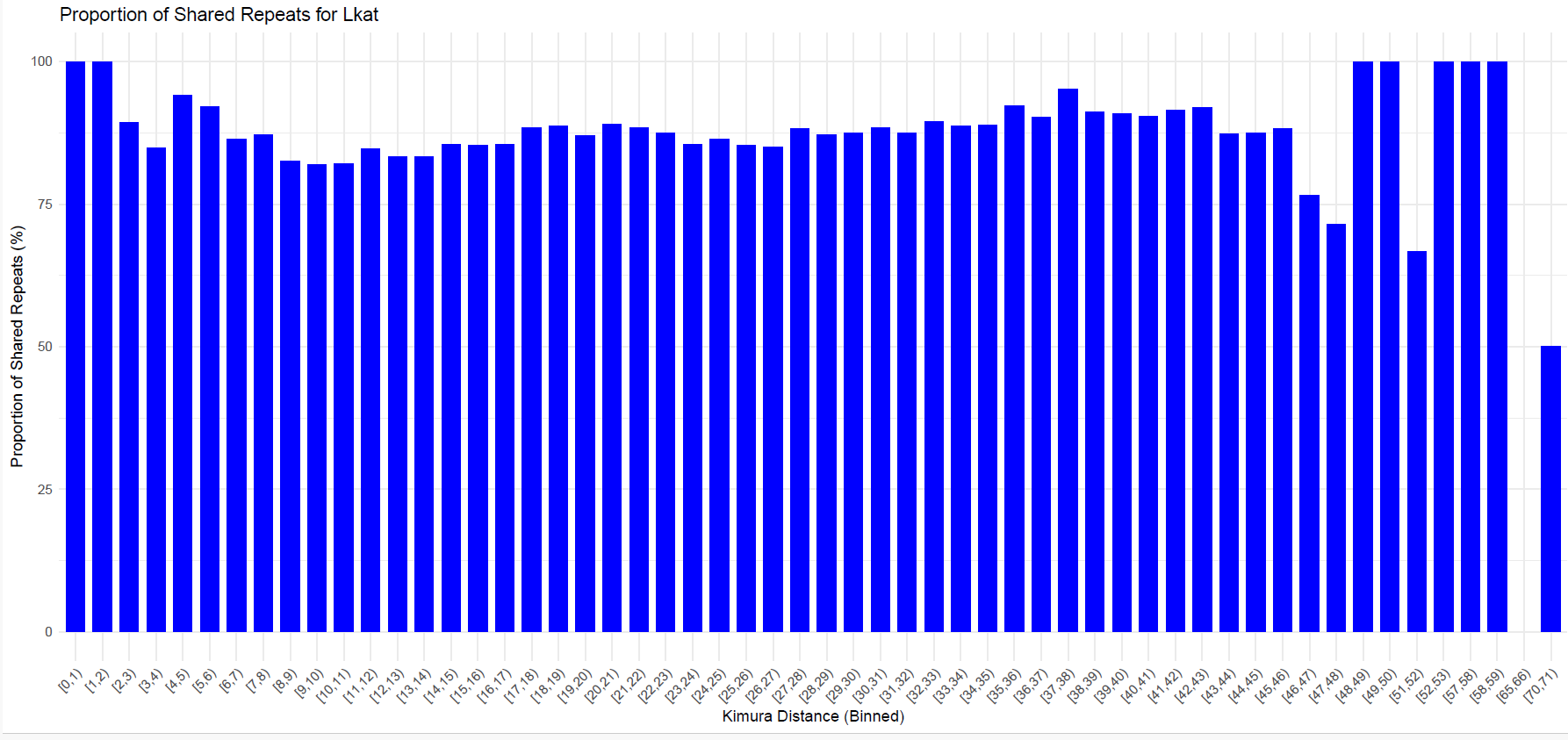

Figure S12. Proportion of shared repeats between the two species in total number of repeats in *L. katipo* arranged by Kimura distance. . One bar represents 1% of the maximum Kimura distance.

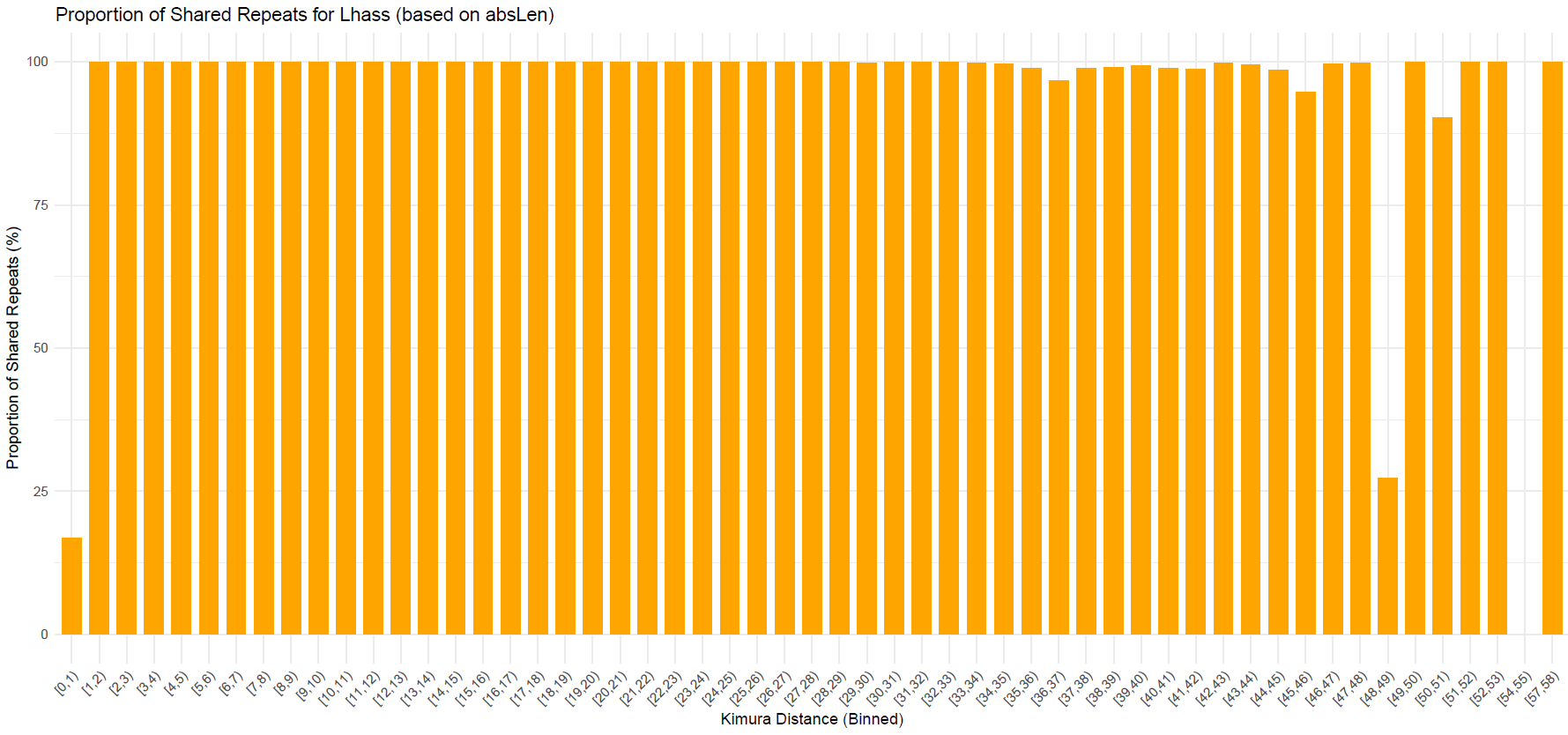

Figure S13. Proportion of total length of shared repeats between the two species in total length of repeats in *L. hasselti* arranged by Kimura distance. . One bar represents 1% of the maximum Kimura distance.

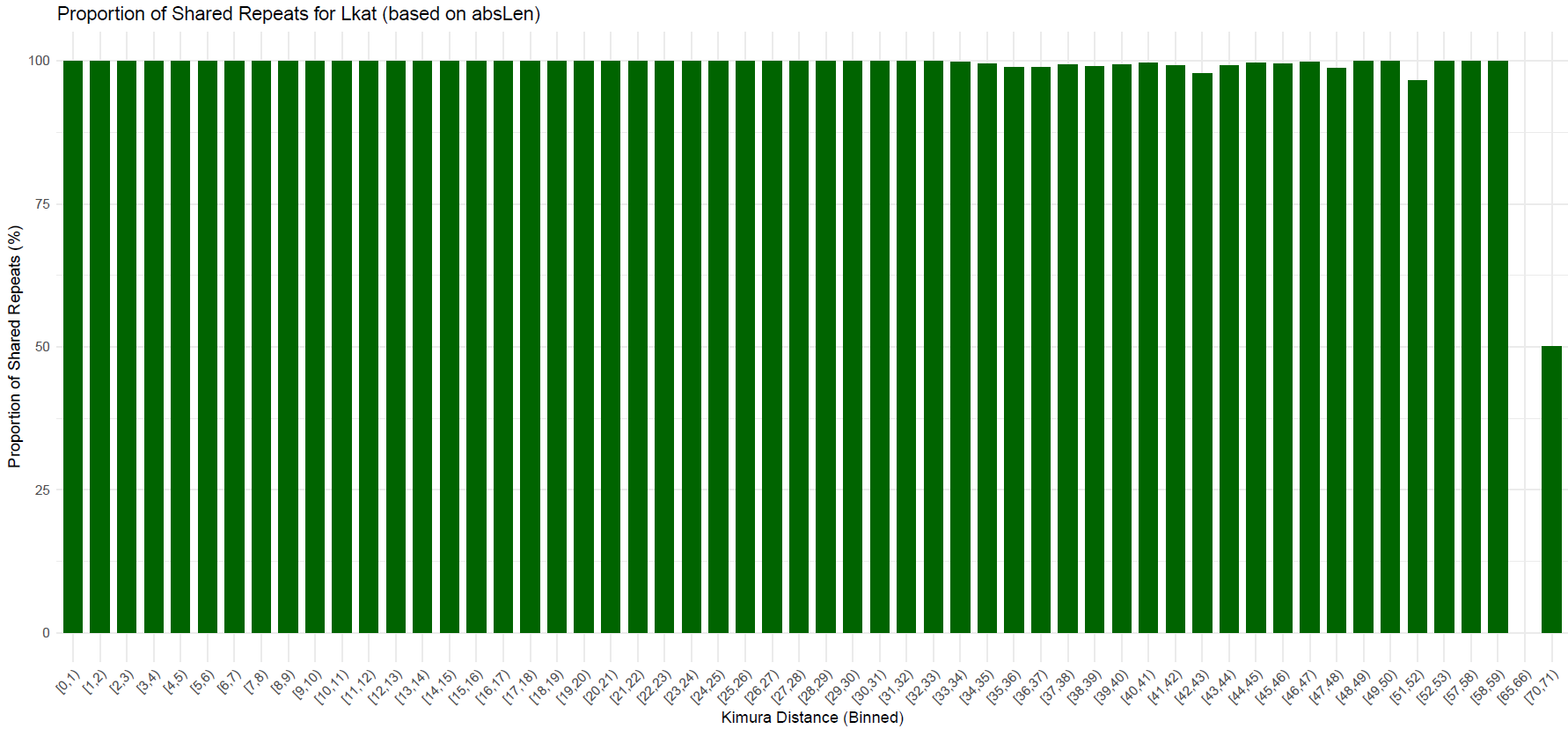

Figure S14. Proportion of total length of shared repeats between the two species in total length of repeats in *L. katipo* arranged by Kimura distance. . One bar represents 1% of the maximum Kimura distance.

### Programs and commands

#############################################

\#	#

\#	Introduction	#

\#	#

#############################################

\## 0. Preface

\# Below are all commands used to assemble genomes of L. katipo and L. hasselti.

\# List of all programs used with versions.

ARIMA pipeline A160156 v02

BLAST v2.13.0 (nt database was downloaded on 24.01.2022)

BlobToolKit v4.4.3

BUSCO v5.5.0

bwa-mem2 v2.2.1

fastp v0.23.4

FastQC v0.12.1

Flye v2.9.2

FigTree v1.4.4

gffread v0.12.8.

HiFiasm v0.19.8

juicer v1.1

juicer_tools v1.22.01

Jupiter plot v1.1

minimap2 v2.26

OrthoFinder version 3.0.1b1

picard v3.1.0-0

Polypolish v0.6.0

Purge_Dups v1.2.6

qualimap v2.3

QUAST v5.2.0

RepeatMasker v4.1.7-p1

RepeatModeler v2.0.5

samtools v1.19.1

TGS-GapCloser v1.2.1

vsearch v2.29.1_linux_x86_64

wtdbg2 v2.5

yahs v1.2a.1.patch

#############################################

\#	#

\#	Latrodectus katipo assembly	#

\#	#

#############################################

######## 1. Raw data QC for CLR, HiC and short read data ########

\# All QC was done using fastqc

\# CLR

fastqc lkati_pacbio_rawdata.fasta.gz --noextract

\# HiC

fastqc Lkati_TAGCTTAT-AGATCTCG_R1.fastq.gz Lkati_TAGCTTAT-AGATCTCG_R2.fastq.gz

\# Short reads

\# cat was used to merge several fastq files from the sequencing run, the coverage is appx. x40 coverage

fastqc LKAT20_merged_1.fq.gz LKAT20_merged_2.fq.gz --noextract

######## 2. CLR assembly. NOTE! BUSCO and QUAST were ran on each intermediate assembly. ########

wtdbg2 -x sq -g 2.5g -t 32 -AS 2 -K 2000 -e 2 -p 16 -k 1 –align-dovetail 2048 -i rawdata.fasta.gz -o Lkatipo_wtdbg2_add_param

wtpoa-cns -t 32 -i Lkatipo_wtdbg2_add_param.ctg.lay.gz -fo Lkatipo_wtdbg2_add_param.ctg.fa

######## 3. Polishing ########

\# 3.1 Long reads polishing

flye --polish-target Lkatipo_wtdbg2_add_param.ctg.fa --pacbio-raw lkati_pacbio_rawdata.fasta.gz -i 3 -t 64 -o . # creates ploished_1.fasta

\# 3.2 Short reads polishing.

\# 3.2.1 Trim short reads.

fastp -i LKAT20_merged.fq.gz -I LKAT20_merged.fq.gz -o LKAT20_merged_trim_1.fq.gz -O LKAT20_merged_trim_1.fq.gz -w 16 --trim_poly_g -j LKAT20_merged_trim.json -h LKAT20_merged_trim.html

\# 3.2.2 Short read polishing commands

\# map F and R reads to the reference

bwa-mem2 mem -t 64 -a polished_1.fasta LKAT20_merged_trim_1.fq.gz > Lkat_alignments_1.sam

bwa-mem2 mem -t 64 -a polished_1.fasta LKAT20_merged_trim_2.fq.gz > Lkat_alignments_2.sam

\# 3.2.3 Polishing

polypolish polish polished_1.fasta Lkat_alignments_1.sam Lkat_alignments_2.sam > Lkatipo_wtdbg2_add_param_polished_long_short.fasta

######## 4. Deduplication/purge ########

\# create config file. clr.txt contains PATH to lkati_pacbio_rawdata.fasta.gz, pbfofn in the manual

pd_config.py -n Lkat_wtdbg2_add_param_short_long_polish_config.json Lkatipo_wtdbg2_add_param_polished_long_short.fasta clr.txt

\# modify config file according to the manual here https://github.com/dfguan/purge_dups

#run purge_dups

run_purge_dups.py Lkat_wtdbg2_add_param_short_long_polish_config.json /PATH_TO_/mambaforge-pypy3/envs/purge_dups/bin/ Lkat_wtdbg2_add_param_short_long_polish -p bash

######## 5. Scaffolding was done using ARIMA mapping pipeline and yahs ########

\# 5.1 Mapping HiC short read data to the reference

\# Index reference genome

samtools faidx Lkatipo_wtdbg2_add_param_polished_long_short.purged.fa

\# Map the short HiC reads. The arima.sh script found here https://github.com/ArimaGenomics/mapping_pipeline/blob/master/arima_mapping_pipeline.sh

\# The arima.sh script

#! /bin/bash

##############################################

\# ARIMA GENOMICS MAPPING PIPELINE 02/08/2019 #

##############################################

#Below find the commands used to map HiC data.

#Replace the variables at the top with the correct paths for the locations of files/programs on your system.

#This bash script will map one paired end HiC dataset (read1 & read2 fastqs). Feel to modify and multiplex as you see fit to work with your volume of samples and system.

##########################################

\# Commands #

##########################################

SRA='Lkati_TAGCTTAT-AGATCTCG'

LABEL='Lkatipo_wtdbg2_add_param_purged_long_short_polish_HiC_katipo'

BWA='/PATH_TO_/mambaforge-pypy3/envs/arima/bin/bwa'

SAMTOOLS='/PATH_TO_/mambaforge-pypy3/envs/arima/bin/samtools'

IN_DIR='/PATH_TO_/DIRECTORY/WITH/fastq/files'

REF='/PATH_TO_/DIRECTORY/WITH/REF_ASSEMBLY/Lkatipo_wtdbg2_add_param_polished_long_short.purged.fa'

FAIDX='$REF.fai'

PREFIX='Lkatipo_wtdbg2_add_param_purged_long_short_polish_HiC_katipo_idx'

RAW_DIR='./bams'

FILT_DIR='./filtered_bams'

FILTER='./filter_five_end.pl'

COMBINER='./two_read_bam_combiner.pl'

STATS='./get_stats.pl'

PICARD='/PATH_TO_/mambaforge-pypy3/envs/arima/share/picard-3.1.0-0/picard.jar'

TMP_DIR='./temp'

PAIR_DIR='paired_bams'

REP_DIR='deduplicated'

REP_LABEL=$LABEL\_rep1

MERGE_DIR='./merged_alignments'

MAPQ_FILTER=10

CPU=32

echo "### Step 0: Check output directories exist & create them as needed"

[ -d $RAW_DIR ] || mkdir -p $RAW_DIR

[ -d $FILT_DIR ] || mkdir -p $FILT_DIR

[ -d $TMP_DIR ] || mkdir -p $TMP_DIR

[ -d $PAIR_DIR ] || mkdir -p $PAIR_DIR

[ -d $REP_DIR ] || mkdir -p $REP_DIR

[ -d $MERGE_DIR ] || mkdir -p $MERGE_DIR

#echo "### Step 0: Index reference" # Run only once! Skip this step if you have already generated BWA index files

$BWA index -a bwtsw -p $PREFIX $REF

echo "### Step 1.A: FASTQ to BAM (1st)"

$BWA mem -t $CPU $PREFIX $IN_DIR/$SRA\_R1.fastq.gz | $SAMTOOLS view -@ $CPU -Sb - > $RAW_DIR/$SRA\_1.bam

echo "### Step 1.B: FASTQ to BAM (2nd)"

$BWA mem -t $CPU $PREFIX $IN_DIR/$SRA\_R2.fastq.gz | $SAMTOOLS view -@ $CPU -Sb - > $RAW_DIR/$SRA\_2.bam

echo "### Step 2.A: Filter 5' end (1st)"

$SAMTOOLS view -h $RAW_DIR/$SRA\_1.bam | perl $FILTER | $SAMTOOLS view -Sb - > $FILT_DIR/$SRA\_1.bam

echo "### Step 2.B: Filter 5' end (2nd)"

$SAMTOOLS view -h $RAW_DIR/$SRA\_2.bam | perl $FILTER | $SAMTOOLS view -Sb - > $FILT_DIR/$SRA\_2.bam

echo "### Step 3A: Pair reads & mapping quality filter"

perl $COMBINER $FILT_DIR/$SRA\_1.bam $FILT_DIR/$SRA\_2.bam $SAMTOOLS $MAPQ_FILTER | $SAMTOOLS view -bS -t $FAIDX - | $SAMTOOLS sort -@ $CPU -o $TMP_DIR/$SRA.bam -

echo "### Step 3.B: Add read group"

java -Xmx4G -Djava.io.tmpdir=temp/ -jar $PICARD AddOrReplaceReadGroups INPUT=$TMP_DIR/$SRA.bam OUTPUT=$PAIR_DIR/$SRA.bam ID=$SRA LB=$SRA SM=$LABEL PL=ILLUMINA PU=none

###############################################################################################################################################################

\###	How to Accommodate Technical Replicates	###

\### This pipeline is currently built for processing a single sample with one read1 and read2 fastq file. ###

\### Technical replicates (eg. one library split across multiple lanes) should be merged before running the MarkDuplicates command. ###

\### If this step is run, the names and locations of input files to subsequent steps will need to be modified in order for subsequent steps to run correctly.###

\### The code below is an example of how to merge technical replicates. ###

###############################################################################################################################################################

\#	REP_NUM=X #number of the technical replicate set e.g. 1

\#	REP_LABEL=$LABEL\_rep$REP_NUM

\#	INPUTS_TECH_REPS=('bash' 'array' 'of' 'bams' 'from' 'replicates') #BAM files you want combined as technical replicates

\#	example bash array - INPUTS_TECH_REPS=('INPUT=A.L1.bam' 'INPUT=A.L2.bam' 'INPUT=A.L3.bam')

\#	java -Xmx8G -Djava.io.tmpdir=temp/ -jar $PICARD MergeSamFiles $INPUTS_TECH_REPS OUTPUT=$TMP_DIR/$REP_LABEL.bam USE_THREADING=TRUE ASSUME_SORTED=TRUE VALIDATION_STRINGENCY=LENIENT

echo "### Step 4: Mark duplicates"

java -Xmx30G -XX:-UseGCOverheadLimit -Djava.io.tmpdir=temp/ -jar $PICARD MarkDuplicates INPUT=$PAIR_DIR/$SRA.bam OUTPUT=$REP_DIR/$REP_LABEL.bam METRICS_FILE=$REP_DIR/metrics.$REP_LABEL.txt TMP_DIR=$TMP_DIR ASSUME_SORTED=TRUE VALIDATION_STRINGENCY=LENIENT REMOVE_DUPLICATES=TRUE

$SAMTOOLS index $REP_DIR/$REP_LABEL.bam

perl $STATS $REP_DIR/$REP_LABEL.bam > $REP_DIR/$REP_LABEL.bam.stats

echo "Finished Mapping Pipeline through Duplicate Removal"

#########################################################################################################################################

\###	How to Accommodate Biological Replicates	###

\### This pipeline is currently built for processing a single sample with one read1 and read2 fastq file. ###

\### Biological replicates (eg. multiple libraries made from the same sample) should be merged before proceeding with subsequent steps.###

\### The code below is an example of how to merge biological replicates. ###

#########################################################################################################################################

\#

\#	INPUTS_BIOLOGICAL_REPS=('bash' 'array' 'of' 'bams' 'from' 'replicates') #BAM files you want combined as biological replicates

\#	example bash array - INPUTS_BIOLOGICAL_REPS=('INPUT=A_rep1.bam' 'INPUT=A_rep2.bam' 'INPUT=A_rep3.bam')

\#

\#	java -Xmx8G -Djava.io.tmpdir=temp/ -jar $PICARD MergeSamFiles $INPUTS_BIOLOGICAL_REPS OUTPUT=$MERGE_DIR/$LABEL.bam USE_THREADING=TRUE ASSUME_SORTED=TRUE VALIDATION_STRINGENCY=LENIENT

\#

\#	$SAMTOOLS index $MERGE_DIR/$LABEL.bam

\# perl $STATS $MERGE_DIR/$LABEL.bam > $MERGE_DIR/$LABEL.bam.stats

\# echo "Finished Mapping Pipeline through merging Biological Replicates"

\# run the script to produce .bam files

./arima.sh

\# 5.2 run yahs and juicer_tools to get .hic, .agp files and the scaffolded fasta is saved under yahs.out_scaffolds_final.fa

yahs Lkatipo_wtdbg2_add_param_polished_long_short.purged.fa Lkatipo_wtdbg2_add_param_purged_long_short_polish_HiC_katipo_rep1.bam

juicer pre -a -o Lkat_wtdbg2_add_param_long_short_polished_purged_Lkat_HiC_out yahs.out.bin yahs.out_scaffolds_final.agp Lkatipo_wtdbg2_add_param_polished_long_short.purged.fa.fai > Lkat_wtdbg2_add_param_long_short_polished_purged_Lkat_HiC_out.log 2>&1

cat Lkat_wtdbg2_add_param_long_short_polished_purged_Lkat_HiC_out.log | grep PRE_C_SIZE | awk '{print $2" "$3}' > chromsize4mancur.txt

java -Xmx150G -jar /PATH_TO_/juicer_tools_1.22.01.jar pre --threads 32 Lkat_wtdbg2_add_param_long_short_polished_purged_Lkat_HiC_out.txt Lkat_wtdbg2_add_param_long_short_polished_purged_Lkat_HiC_out.hic.part chromsize4mancur.txt

mv Lkat_wtdbg2_add_param_long_short_polished_purged_Lkat_HiC_out.hic.part Lkat_wtdbg2_add_param_long_short_polished_purged_Lkat_HiC_out.hic

######## 6. Gap closing ########

cat lkati_pacbio_rawdata.fasta.gz | tgsgapcloser --ne --tgstype pb --minmap_arg " -x asm20" --scaff yahs.out_scaffolds_final.fa --reads /dev/fd/0 --output Lkatipo_fin_scaffolded_no_err_corr_002 --thread 90 >pipe.log 2>pipe.err

######## 7. Remove contamination and visualize using blobtoolkit ########

\# The pipeline also requires:

\# BUSCO genes, in this case arachnida_odb10

\# NCBI nt database, download date is 24.01.2022

\# 7.1 Map the raw CLR reads against the fasta obtained in step 6 and sort and index .bam.

minimap2 -t 32 -a -t 32 -o Lkatipo_scaff_gaps.sam Lkatipo_fin_scaffolded_no_err_corr_002.scaff_seqs lkati_pacbio_rawdata.fasta.gz

samtools sort -O bam -@32 -o Lkatipo_scaff_gasps_sorted.bam Lkatipo_scaff_gaps.sam

samtools index -@ 24 -c Lkatipo_scaff_gasps_sorted.bam Lkatipo_scaff_gaps.bam

\# 7.2 BLAST the fasta obtained in step 6 against NCBI nt database. Access date 24.01.2022

nice -n 5 blastn -task megablast \

-query Lkatipo_fin_scaffolded_no_err_corr_002.scaff_seqs \

-db /PATH_TO_/BLAST_db/nt \

-max_target_seqs 1 -max_hsps 1 \

-outfmt '6 qseqid staxids bitscore std' \

-num_threads 64 -evalue 1e-25 -out ./Lkatipo_scaff_gaps.blast

\# 7.3 Run the blobtools pipeline

blobtools create --threads 32 --fasta Lkatipo_fin_scaffolded_no_err_corr_002.scaff_seqs --create blobtoolsdir --cov Lkatipo_scaff_gaps_shortread_sorted.bam --cov Lkatipo_scaff_gasps_sorted.bam

blobtools add --threads 64 --hits Lkatipo_scaff_gaps.blast --taxrule bestsumorder --taxdump /PATH_TO_/taxdump_240122 blobtoolsdir

blobtools add --threads 32 --busco /PATH_TO_/run_arachnida_odb10/full_table.tsv blobtoolsdir

blobtk plot -d blobtoolsdir -v blob -o Lkatipo_fin_scaffolded_no_err_corr_002.svg

blobtk plot -d blobtoolsdir -v cumulative -o Lkatipo_fin_scaffolded_no_err_corr_002_cumulative.svg

blobtk plot -d blobtoolsdir -v snail -o Lkatipo_fin_scaffolded_no_err_corr_002_snail.svg

\## Filtering. First create a list of taxa without the ones that were excluded

\# The command to list all phyla with the hit

blobtools view -i output_directory/assembly.fasta.blobDB.json -T phylum -o phylum_hits.txt

blobtools filter --fasta Lkatipo_fin_scaffolded_no_err_corr_002.scaff_seqs --output "/PATH_TO_OUTPUT_DIRECTORY/" --param length--Min=1000 --param bestsumorder_phylum--Keys=Chlamydiota,Pseudomonadota,Mollusca,Mycoplasmatota,Chordata,Uroviricota,Cnidaria --param Lkatipo_scaff_gaps_shortread_sorted_cov--Min=3 --param Lkatipo_scaff_gasps_sorted_cov--Min=3 blobtoolsdir

\# after that use custome script to filter the original fasta.

python filter_script.py

\## filter_script.py is below

import json

def filter_fasta(original_fasta, identifiers_file, output_fasta):

\# Load contig names from identifiers.json

with open(identifiers_file, 'r') as identifiers_json:

identifiers_data = json.load(identifiers_json)

contig_names_to_keep = set(identifiers_data["values"])

\# Filter original fasta and write to output fasta

with open(original_fasta, 'r') as original, open(output_fasta, 'w') as output:

write_sequence = False

for line in original:

if line.startswith(">"):

header = line.strip()[1:] # Extract the header without ">"

write_sequence = header in contig_names_to_keep

if write_sequence:

output.write(line)

\# Usage

original_fasta = "/PATH_TO_/Lkatipo_fin_scaffolded_no_err_corr_002.scaff_seqs"

identifiers_file = "/PATH_TO_/identifiers.json"

output_fasta = "Lkatipo_fin_filtered.fasta"

filter_fasta(original_fasta, identifiers_file, output_fasta)

######## 8. Assembly QC ########

\# 8.1 BUSCO

busco -i "Lkatipo_fin_filtered.fasta" -o "Lkatipo_fin_filtered_busco" -m genome -l arachnida_odb10 -c 64

\# 8.2 QUAST

quast -o Lkatipo_fin_filtered_QUAST -t 32 Lkatipo_fin_filtered.fasta

######## 9. Mapping two L. katipo males against the final genome assembly ########

bwa-mem2 mem Male1_R1.fastq.gz Male1_R2.fastq.gz | samtools view -b -@ 4 - | samtools sort -@ 4 -o Male1.bam

qualimap bamqc -bam Male1.bam -nt 16 -c -outdir /OUTPUT_DIR/ --java-mem-size=64G -outformat PDF

#############################################

\#	#

\#	Latrodectus hasselti assembly	#

\#	#

#############################################

```
## NOTE! The assembly largely followed the same steps as L. katipo that's why only steps different from the listed above are listed.
## NOTE! Since there was no HiC data and short reads for polishing were not necessary, as well as polishing step with HiFi data, steps 3 and 5 were not done.
## NOTE! BUSCO and QUAST were ran on each intermediate assembly.
```

######## 1. Raw input data manipulation and QC ########

\# Conversion of CSC reads to fastq

samtools fastq input.bam > input.fastq

gzip input.fastq

\# QC

fastqc input.fastq.gz --noextract

######## 2. Assembly with HiFiasm ########

hifiasm -o hifiasm_output.asm -t 32 input.fastq.gz

```
## The assembly continued with steps: 4. Deduplication/purge; 6. Gap closing; 7. Remove contamination and visualize using blobtoolkit; 8. Assembly QC
## Decontamination steps detected different groups of organisms: Pseudomonadota, Chordata
## 10. Jupiter plot of L. hasselti contigs against L. katipo
```

jupiter name=Lhasselti_circos_to_Lkatipo_13chr_ng90 ref="/PATH_TO_/Lkatipo_fin_scaff_filtered_001_13chrom.fasta" fa="/PATH_TO_//Lhasselti_default_filtered_noPseudChord.purged.fa" sam="/PATH_TO_/Lhasselti_circos_to_Lkatipo_13chr-agp.sam" ng=90 labels=both

#################################

\#	#

\#	Repeats analysis	#

\#	#

#################################

######## 11. RepeatModeler to identify repeats in respective genomes ########

\# 11.1 Build database out of the genome

BuildDatabase -name Lkatipo_repeats_db -engine ncbi Lkatipo_fin_scaff_filtered_001_13chrom.fasta

\# 11.2 Run RepeatModeler. If we ran out of time we added -recoverDir option

RepeatModeler -threads 24 -engine ncbi -database Lkatipo_repeats_db 2>&1 | tee 00_repeatmodeler_Lkatipo_repeats_13chr.log

######## 12. RepeatMasking ########

\# 12.1 Combine identified repeat sequences from both genomes and invertebrate RepBase

cat L.katipo_repdb-families.fa Lhasselti_repdb-familieis.fa invrep.ref > merged_replib.fa

\# 12.2 Run vsearch to sort the sequences and remove duplicates

vsearch --sortbylength merged_replib.fa --output merged_replib_sorted.fa --log vsearch.log

vsearch --cluster_fast merged_replib_sorted.fa --id 0.95 --minseqlength 0 --centroids dedup_replib_centroids.fa --uc dedup_replib_results.uc --consout dedup_replib_consensus.fa --msaout dedup_replib_aligned.fa --log dedup_vsearch.log --threads 24

\# 12.3 Run RepeatMasker to collect repeats statistics and mask the reference genomes

RepeatMasker -pa 24 -gccal -nocut -s -xsmall -a -gff -lib dedup_merged_replib_inv_latr_sorted_consensus.fa -dir /OUTUPUT/DIR/ INPUT_GENOME.fasta

######## 13. Plotting the output ########

\# 13.1 RepeatMasker output requires re-formatting before plotting. Below is the script to do exactly that.

\# This script takes in a RepeatMasker directory and generates the repeat landscape of the contained genome.

\# The script expects a directory named after the species as input

repeatmasker_dir=/PATH_TO_/REPEATMASKER/FILES/miniforge3/envs/repeats/share/RepeatMasker

export PERL5LIB=$repeatmasker_dir

indir=$1

pref=$(basename $indir)

g_size=$(cat $indir/*.tbl | grep "total length:" | awk '{print $3}') # | rev | cut -c4- | rev)

echo "Indir: ${indir}"

echo "Prefix: ${pref}"

echo "Genomesize: ${g_size}"

$repeatmasker_dir/util/calcDivergenceFromAlign.pl -s $indir/$pref".divsum" $indir/*.align

$repeatmasker_dir/util/createRepeatLandscape.pl -div $indir/$pref".divsum" -g $g_size > $pref"_repeatlandscape.html"

\# The .divsum file contains the matrix in the end that is used for plotting. Below is a sample of the matrix.

Coverage for each repeat class and divergence (Kimura)

Div DNA/Academ-1 DNA/CMC-Chapaev-3 DNA/CMC-EnSpm DNA/Ginger-2 DNA/Ginger-2 DNA/Kolobok-E DNA/Kolobok-Hydra DNA/Kolobok-Hydra DNA/MULE-MuDR DNA/MULE-MuDR DNA/Maverick DNA/Merlin DNA/Merlin DNA/P DNA/P DNA/P DNA/PIF-Harbinger DNA/PIF-Harbinger DNA/PIF-ISL2EU DNA/PiggyBac DNA/PiggyBac DNA/Sola-1 DNA/Sola-2 DNA/TcMar-Fot1 DNA/TcMar-Mariner DNA/TcMar-Mariner DNA/TcMar-Mariner DNA/TcMar-Mariner DNA/TcMar-Pogo DNA/TcMar-Pogo DNA/TcMar-Pogo DNA/TcMar-Tc1 DNA/TcMar-Tc1 DNA/TcMar-Tc1 DNA/TcMar-Tc1 DNA/TcMar-Tc1 DNA/TcMar-Tc2 DNA/TcMar-Tc2 DNA/TcMar-Tigger DNA/TcMar-Tigger DNA/TcMar-Tigger DNA/hAT-Blackjack DNA/hAT-Blackjack DNA/hAT-Blackjack DNA/hAT-Charlie DNA/hAT-Charlie DNA/hAT-Charlie DNA/hAT-Charlie DNA/hAT-Charlie DNA/hAT-Pegasus DNA/hAT-Tip100 DNA/hAT-Tip100 DNA/hAT-Tip100 DNA/hAT-hAT5 DNA/hAT-hAT5 DNA/hAT-hAT5 DNA/hAT-hATx DNA LINE/I-Jockey LINE/I-Jockey LINE/I-Jockey LINE/I-Jockey LINE/I LINE/I LINE/I LINE/I LINE/Penelope LINE/R1-LOA LINE/R1-LOA LINE/R1-LOA LINE/R1 LINE/R1 LINE/R1 LINE/RTE-BovB LINE/RTE-BovB LINE/RTE-BovB LINE/RTE LINE/Tad1 LINE LINE LTR/Copia LTR/Copia LTR/Gypsy LTR/Gypsy LTR/Gypsy LTR/Gypsy LTR/Pao LTR/Pao RC/Helitron Simple_repeat Simple_repeat Unknown Unknown Unknown Unknown Unspecified tRNA

0 0 0 3461 103 124 223 78294 0 66574 2069 530618 246 0 6339 7487 767 85835 0 75 234638 0 595 124424 42635 335936 123873 317 460 32 0 444 202443 22399 302 6279 122 0 260 123 191 108 403 0 540 14597 10822 1455 6233 0 0 3217 0 71 71 125 0 0 2175 81250 2243 1027 188 2946 80366 58 840 2223 1803 7203 71 6778 933 976 6880 105 15140 759 0 5638 283 1154 53262 2073283 359438 103478 81184 383963 170 764 0 0 1001145 344333 215248 323 31333 17544

1 653 53 41375 384 331 1380 1543 0 128653 65428 364119 490 0 116055 199606 78110 93822 0 9515 530752 138 740 25921 31419 5675 227500 107 0 131 0 12725 3491031 690710 223 48940 0 0 0 56 144 295 1113 60348 306 238781 113294 26091 323252 0 0 17231 447 70 52 5750 0 115 8488 347115 257677 10266 208 23516 912449 3227 912 38070 3581 381706 437 121926 84370 59091 22605 229 144396 479 0 46991 253 5391 362442 2005072 2186397 52984 114430 281622 250349 0 0 0 1685990 1512311 112042 336 37140 50336

2 53890 0 153752 1412 898 3265 98 0 180257 254819 179104 1111 179 88215 275204 309596 95828 445 80423 112368 321 6394 50794 15402 10355 139882 632 1166 128 0 54730 3746683 4196241 4678 97841 0 35 382 228 486 119 11870 354754 982 335852 428343 257486 557202 990 0 77281 2650 369 39 52098 0 42 60243 859080 296141 59440 5773 84923 683275 66814 6300 358090 14265 1354876 3488 276840 83729 362233 45350 4354 189465 4805 0 182751 366 4912 69717 990737 1068830 147950 50497 416922 179718 2525 0 0 3332346 1828361 35483 67481 69427 55746

\# simple copying to the new file and piping into R is enough, however, in our case we had to add additional column named Unassigned so the file can be read correctly.

\# Also the script added unnecessary seqs counts like DNA/Academ-1;seqs=1 that were removed with sed, e.g.

sed 's/;seqs=[0-9]*//g' "002_softmasked_hasselti.divsum" > 002_softmasked_corr.divsum

\# 13.2 R script to plot the matrix of repetative elements

\# packages needed

library(reshape2)

library(dplyr)

library(ggplot2)

library(viridisLite)

library(viridis)

library(hrbrthemes)

library(tidyverse)

library(gridExtra)

#attached base packages:

\# [1] stats	graphics grDevices utils	datasets methods	base

#other attached packages:

\# [1] gridExtra_2.3	forcats_0.5.0	stringr_1.4.0	dplyr_0.8.5	purrr_0.3.4

\# [6] readr_1.3.1	tidyr_1.1.0	tibble_3.0.1	tidyverse_1.3.0	hrbrthemes_0.8.0

\# [11] viridis_0.5.1	viridisLite_0.3.0 ggplot2_3.3.0	reshape_0.8.8

\# Read the file with the matrix

KimuraDistance <- read.csv("COPIED_MATRIX.txt",sep="", row.names=NULL)

\# Remove unnecessary column

KimuraDistance <- KimuraDistance %>% select(-Unassigned)

#add here the genome size in bp

genomes_size=1372912898

\# Calculate the Kimura distances and the percent of the genome covered by each family representative of each distance bin

kd_melt = melt(KimuraDistance, id="Div")

kd_melt$norm = kd_melt$value/genomes_size * 100

kd_melt$Class = unlist(lapply(strsplit(as.character(kd_melt$variable), "\\."), '[[', 1))

kd_melt <- kd_melt %>%

group_by(Div, Class) %>%

summarise(

value = sum(value, na.rm = TRUE),

norm = sum(norm, na.rm = TRUE),

.groups = "drop"

)

\# plot the resulting matrix. It is expected that the shorter the Kimura distance the more active the repeats are.

ggplot(kd_melt, aes(fill=Class, y=norm, x=Div)) +

geom_bar(position="stack", stat="identity",color="black") +

scale_fill_viridis(discrete = T, option="turbo", begin = 0.1, direction=1) +

theme_classic() +

xlab("Kimura substitution level") +

ggtitle("PLOT TITLE") +

ylab("Percent of the genome") +

labs(fill = "") +

coord_cartesian(xlim = c(0, 55)) +

theme(axis.text=element_text(size=11),axis.title =element_text(size=12))

\# 13.3 R script to count and plot overlapping repeats between the species

library(dplyr)

library(ggplot2)

file1 <- "Lkat_ALL_Kimura_dist_for_barchart_plotting.txt"

file2 <- "Lhas_Kimura_dist_for_barchart_plotting.txt"

\# Read the files

data1 <- read.delim(file1, header = TRUE, sep = "\t")

data2 <- read.delim(file2, header = TRUE, sep = "\t")

\# Add an identifier column for each file

data1$ID <- "katipo"

data2$ID <- "hasselti"

data1 <- filter(data1, absLen > 0, wellCharLen > 0, Kimura. > 0)

data2 <- filter(data2, absLen > 0, wellCharLen > 0, Kimura. > 0)

\# Find common 'Repeat' values

common_repeats <- intersect(data1$Repeat, data2$Repeat)

\# Extract matching rows from both files

filtered1 <- filter(data1, Repeat %in% common_repeats)

filtered2 <- filter(data2, Repeat %in% common_repeats)

\# Combine the extracted data

combined_data <- bind_rows(filtered1, filtered2)

\# Write to output file

write.table(combined_data, "matched_repeats_Lhas_Lkat_ALL.txt", sep = "\t", row.names = FALSE, quote = FALSE)

\# Define file paths

#file_lhass <- "Lhas_Kimura_dist_for_barchart_plotting.txt" # Original Lhass file

#file_lkat <- "Lkat_Kimura_dist_for_barchart_plotting.txt"	# Original Lkat file

#file_merged <- "matched_repeats_Lhas_Lkat.txt" # Merged file with labeled repeats

\# Read the files

#data_lhass <- read.delim(file_lhass, header = TRUE, sep = "\t")

#data_lkat <- read.delim(file_lkat, header = TRUE, sep = "\t")

#data_merged <- read.delim(file_merged, header = TRUE, sep = "\t")

\# Filter for positive values in 'absLen' and 'wellCharLen'

#data_lhass <- filter(data_lhass, absLen > 0, wellCharLen > 0, Kimura. > 0)

#data_lkat <- filter(data_lkat, absLen > 0, wellCharLen > 0, Kimura. > 0)

#data_merged <- filter(data_merged, absLen > 0, wellCharLen > 0, Kimura. > 0)

\# Filter merged data to only count Repeat values with matching IDs

shared_lhass <- filter(combined_data, ID == "hasselti")

shared_lkat <- filter(combined_data, ID == "katipo")

\# Define bins from 0 to 60 with a step of 1 (i.e., 0-1, 1-2, ..., 59-60)

bins <- seq(0, 75, by = 1)

\# Function to compute proportions in bins

compute_proportions <- function(shared_data, original_data, bins) {

\# Count shared repeats in the merged file

shared_binned <- shared_data %>%

mutate(bin = cut(Kimura., breaks = bins, include.lowest = TRUE, right = FALSE)) %>%

group_by(bin) %>%

summarise(shared_count = n(), .groups = "drop")

\# Count total repeats in the original dataset (file1.txt or file2.txt) per bin

original_binned <- original_data %>%

mutate(bin = cut(Kimura., breaks = bins, include.lowest = TRUE, right = FALSE)) %>%

group_by(bin) %>%

summarise(total_count = n(), .groups = "drop")

\# Merge the shared and original counts by bin

binned_data <- full_join(shared_binned, original_binned, by = "bin") %>%

mutate(proportion = (shared_count / total_count) * 100) # Calculate proportion

return(binned_data)

}

\# Compute proportions for Lhass and Lkat

lhass_prop <- compute_proportions(shared_lhass, data2, bins)

lkat_prop <- compute_proportions(shared_lkat, data1, bins)

\# Plot for Lhass

p1 <- ggplot(lhass_prop, aes(x = bin, y = proportion, fill = "Lhass")) +

geom_bar(stat = "identity", width = 0.7, fill = "red") +

theme_minimal() +

labs(title = "Proportion of Shared Repeats for Lhass",

x = "Kimura Distance (Binned)",

y = "Proportion of Shared Repeats (%)") +

theme(axis.text.x = element_text(angle = 45, hjust = 1)) +

scale_fill_manual(values = c("Lhass" = "red"), guide = "none")

\# Plot for Lkat

p2 <- ggplot(lkat_prop, aes(x = bin, y = proportion, fill = "Lkat")) +

geom_bar(stat = "identity", width = 0.7, fill = "blue") +

theme_minimal() +

labs(title = "Proportion of Shared Repeats for Lkat",

x = "Kimura Distance (Binned)",

y = "Proportion of Shared Repeats (%)") +

theme(axis.text.x = element_text(angle = 45, hjust = 1)) +

scale_fill_manual(values = c("Lkat" = "blue"), guide = "none")

\# Display the plots

print(p1)

print(p2)

\# Same for the absLength

\# Function to compute proportions based on absLen in bins

compute_proportions <- function(shared_data, original_data, bins) {

\# Calculate the total shared length in each bin (from merged data)

shared_binned <- shared_data %>%

mutate(bin = cut(Kimura., breaks = bins, include.lowest = TRUE, right = FALSE)) %>%

group_by(bin) %>%

summarise(shared_length = sum(absLen), .groups = "drop")

\# Calculate the total length of repeats in the original dataset (file1.txt or file2.txt) per bin

original_binned <- original_data %>%

mutate(bin = cut(Kimura., breaks = bins, include.lowest = TRUE, right = FALSE)) %>%

group_by(bin) %>%

summarise(total_length = sum(absLen), .groups = "drop")

\# Merge the shared and original lengths by bin

binned_data <- full_join(shared_binned, original_binned, by = "bin") %>%

mutate(proportion = (shared_length / total_length) * 100) # Calculate proportion based on lengths

return(binned_data)

}

\# Compute proportions for Lhass and Lkat based on absLen

lhass_prop <- compute_proportions(shared_lhass, data2, bins)

lkat_prop <- compute_proportions(shared_lkat, data1, bins)

\# Plot for Lhass

p1 <- ggplot(lhass_prop, aes(x = bin, y = proportion, fill = "Lhass")) +

geom_bar(stat = "identity", width = 0.7, fill = "orange") +

theme_minimal() +

labs(title = "Proportion of Shared Repeats for Lhass (based on absLen)",

x = "Kimura Distance (Binned)",

y = "Proportion of Shared Repeats (%)") +

theme(axis.text.x = element_text(angle = 45, hjust = 1)) +

scale_fill_manual(values = c("Lhass" = "orange"), guide = "none")

\# Plot for Lkat

p2 <- ggplot(lkat_prop, aes(x = bin, y = proportion, fill = "Lkat")) +

geom_bar(stat = "identity", width = 0.7, fill = "darkgreen") +

theme_minimal() +

labs(title = "Proportion of Shared Repeats for Lkat (based on absLen)",

x = "Kimura Distance (Binned)",

y = "Proportion of Shared Repeats (%)") +

theme(axis.text.x = element_text(angle = 45, hjust = 1)) +

scale_fill_manual(values = c("Lkat" = "darkgreen"), guide = "none")

\# Display the plots

print(p1)

print(p2)

\## PLOT SCATTER AND t-test

katipo_group <- filter(combined_data, ID == "katipo")

hasselti_group <- filter(combined_data, ID == "hasselti")

missing_in_hasselti <- setdiff(katipo_group$Repeat, hasselti_group$Repeat)

missing_in_katipo <- setdiff(hasselti_group$Repeat, katipo_group$Repeat)

\# Print the results

if (length(missing_in_hasselti) > 0) {

cat("Missing in hasselti:\n", missing_in_hasselti, "\n")

} else {

cat("No missing values in hasselti.\n")

}

if (length(missing_in_katipo) > 0) {

cat("Missing in katipo:\n", missing_in_katipo, "\n")

} else {

cat("No missing values in katipo.\n")

}

\# Sort the katipo group by Kimura values (ascending order)

katipo_group <- katipo_group %>% arrange(Kimura.)

katipo_group_leng <- katipo_group %>% arrange(absLen)

\# Reorder the hasselti group to match the order of katipo based on the Repeat column

hasselti_group <- hasselti_group %>%

slice(match(katipo_group$Repeat, Repeat))

hasselti_group_leng <- hasselti_group %>%

slice(match(katipo_group_leng$Repeat, Repeat))

\# Create the scatter plot

p <- ggplot() +

geom_point(aes(x = katipo_group$Kimura., y = hasselti_group$Kimura.), color = "#1f77b4") +

theme_minimal() +

labs(title = "Comparison of Kimura Distance (Katipo vs Hasselti)",

x = "Kimura Distance (Katipo)",

y = "Kimura Distance (Hasselti)")

\# Display the plot

print(p)

length1 <- ggplot() +

geom_point(aes(x = katipo_group_leng$absLen, y = hasselti_group_leng$absLen), color = "red") +

theme_minimal() +

labs(title = "Comparison of repeat absolute length (Katipo vs Hasselti)",

x = "absLength Distance (Katipo)",

y = "absLength Distance (Hasselti)")

length1

t.test(katipo_group$Kimura., y = hasselti_group$Kimura.)

#############################

Annotation

#############################

14. Braker command

BRAKER CALL: /opt/BRAKER/scripts/braker.pl --species=hasselti --AUGUSTUS_CONFIG_PATH=/scratch/utr_ivanov/002_Lhasselti_genome/005_annotation_Lhasselti_2/config/ --genome=Lhasselti_default_filtered_purged_reordered.fa.masked --prot_seq=Arthropoda.fa --rnaseq_sets_ids=HPOOL1,HPOOL2,AFRB13,AMRB25,PFRB15,PMSRB53 --rnaseq_sets_dirs=/scratch/utr_ivanov/002_Lhasselti_genome/005_annotation_Lhasselti/000_RNAseq_hasselti/ --workingdir=001_Lhass_annot_Arthrdb --threads 24 --busco_lineage arachnida_odb10

15. eggNOG command (run on the website)

emapper.py --cpu 20 --mp_start_method forkserver --data_dir /dev/shm/ -o out --output_dir /emapper_web_jobs/emapper_jobs/user_data/MM_2esidf11 --temp_dir /emapper_web_jobs/emapper_jobs/user_data/MM_2esidf11 --override -m diamond --dmnd_ignore_warnings -i /emapper_web_jobs/emapper_jobs/user_data/MM_2esidf11/queries.fasta --evalue 0.001 --score 60 --pident 40 --query_cover 20 --subject_cover 20 --itype proteins --tax_scope 6656 --target_orthologs all --go_evidence non-electronic --pfam_realign none --report_orthologs --decorate_gff yes --excel > /emapper_web_jobs/emapper_jobs/user_data/MM_2esidf11/emapper.out 2> /emapper_web_jobs/emapper_jobs/user_data/MM_2esidf11/emapper.err

16. BUSCO for amino acid sequences

busco -i braker.aa -o Lhasselti_protein_BUSCO -m proteins -l arachnida_odb10 -c 32 -f

#############################

Orthofinder

#############################

\## We investigated the orthogroups for several species in order to identfy genes unique for L. hasselti and L. katipo

\# First, we selected genes and transcripts of L. hasselti that were found only on the contigs mapped to L. katipo

\# Then we selected the longest transcripts (if applicable) for each species.

16. Selection of the longest contig.

sed -e 's/\r//g' -e '/^$/d' Latrodectus_hasselti.faa > cleaned.faa

awk '

/^>/ {

if (seq != "") {

\# print previous record

print header "\t" seq

}

header = $0

seq = ""

next

}

{ seq = seq $0 }

END {

if (seq != "") print header "\t" seq

}

' cleaned.faa |

awk -F'\t' '

{

hdr = $1

seq = $2

gsub(/\*$/, "", seq) # remove trailing *

id = hdr

sub(/^>/, "", id)

split(id, parts, "\\.t") # split at ".t" literally

gene = parts[1]

len = length(seq)

if (len > bestlen[gene]) {

bestlen[gene] = len

besthdr[gene] = hdr

bestseq[gene] = seq

}

}

END {

for (g in besthdr) {

print besthdr[g]

print bestseq[g]

}

}' > proteins_longest.faa

\## Then I extracted names of genes with

grep ">" file.faa > 000_list_of_gene_names.txt

\# and compared to contigs that were mapped to L. katipo. All genes belonged to mapped contigs

\# In order to provide broader context we selected 9 spider species with annotated genomes and available .faa files.

\## List of included taxa:

\# Outgroup non-Theridiidae with aa

GCF_947563725.1	qqArgBrue1.1	Araneidae	Argiope bruennichi	https://ftp.ncbi.nlm.nih.gov/genomes/all/GCF/947/563/725/GCF_947563725.1_qqArgBrue1.1/GCF_947563725.1_qqArgBrue1.1_protein.faa.gz

GCA_019974015.1	Npil_1.0	Araneidae 	Nephila pilipes	https://ftp.ncbi.nlm.nih.gov/genomes/all/GCA/019/974/015/GCA_019974015.1_Npil_1.0/GCA_019974015.1_Npil_1.0_protein.faa.gz

GCF_010614865.2	ASM1061486v2	Eresidae 	Stegodyphus dumicola	https://ftp.ncbi.nlm.nih.gov/genomes/all/GCF/010/614/865/GCF_010614865.2_ASM1061486v2/GCF_010614865.2_ASM1061486v2_protein.faa.gz

GCA_019343175.1	Ogib_1.0	Linyphiidae	Oedothorax gibbosus	https://ftp.ncbi.nlm.nih.gov/genomes/all/GCA/019/343/175/GCA_019343175.1_Ogib_1.0/GCA_019343175.1_Ogib_1.0_protein.faa.gz

\# Outgroup non-Theridiidae with gbff

GCA_051201355.1	IOZ_Pimo_1.0	Pimoidae	Pimoa clavata	Got faa from here https://www.scidb.cn/en/detail?dataSetId=253c4c2a54114b3699e791cbd7a0169c

GCA_947070885.1	tetragnatha_kauaiensis_v1	Tetragnathidae	Tetragnatha kauaiensis	https://datadryad.org/downloads/file_stream/1158590

\# Outgroup Theridiidae

GCA_030067965.1	ASM3006796v1	Theridiidae	Latrodectus elegans	https://s3.ap-northeast-1.wasabisys.com/gigadb-datasets/live/pub/10.5524/102001_103000/102210/Latrodectus_elegans_EVM.out.gff3.pep.all.function

GCA_037975125.2	ASM3797512v2	Theridiidae	Latrodectus hesperus	The aa was obtained externally, https://agdatacommons.nal.usda.gov/articles/dataset/Latrodectus_hesperus_genome_annotations_v0_5_3/24662235?file=43633890

GCF_043381705.1	CAS_Ptep_4.0	Theridiidae	Parasteatoda tepidariorum	https://ftp.ncbi.nlm.nih.gov/genomes/all/GCF/043/381/705/GCF_043381705.1_CAS_Ptep_4.0/GCF_043381705.1_CAS_Ptep_4.0_protein.faa.gz

\# Ingroup

JBNCAM000000000	NA	Theridiidae	Latrodectus katipo	Locla file,

JBMHEB000000000	NA	Theridiidae	Latrodectus hasselti	Local file,

17. Creating a faa file for the L. elegans since it had only gff3 file. We used reference genome and gffread

gffread Latrodectus_elegans_EVM.out.gff3 -g Latrodectus_elegans_genome_sm.fasta -y Latrodectus_elegans.faa -w Latrodectus_elegans_transcripts.fna

\# For the rest of the species we have located the faa files.

18. Orthofinder

\# The dots in the sequences and the end of the line are a problem for diamond.

\# It can also concern end of the line

\# solution is to remove dots both from seqeunces and the end of the line

\# tools are sed (found here https://github.com/davidemms/OrthoFinder/issues/175)

sed -i -e ':a;N;$!ba;s/\.\n/\n/g' 000_input_aa/*.faa

\# and awk

for f in 000_input_aa/*.faa; do awk '/^>/ {print; next} {gsub(/\./, ""); print}' "$f" > "${f%.faa}_clean.faa"; done

\# then the orthofinder was called with:

orthofinder -t 28 -a 28 -f /PATH_TO_THE/cleaned_input_aa/

\# After the orthofinder finished we visalised the trees with FigTree v1.4.4
